# The effect of MurM and a branched cell wall structure on penicillin resistance in *Streptococcus pneumoniae*

**DOI:** 10.1101/2025.04.05.647355

**Authors:** Ragnhild Sødal Gjennestad, Maria Victoria Heggenhougen, Anja Ruud Winther, Johanne Moldstad, Vegard Eldholm, Morten Kjos, Leiv Sigve Håvarstein, Daniel Straume

**Affiliations:** Faculty of Chemistry, Biotechnology and Food Science, Norwegian University of Life Sciences, Ås, Norway; Department of Bacteriology, Norwegian Institute of Public Health, Oslo, Norway

## Abstract

The aminoacyltransferase MurM is an important penicillin resistance determinant in *Streptococcus pneumoniae*. This enzyme attaches a serine or alanine to the dipeptide side chain of lipid II, resulting in branched muropeptides that can be crosslinked to stem peptides in the peptidoglycan layer by penicillin binding proteins (PBPs). Deletion of *murM* results in only linear muropeptides, and more importantly a significant reduction in resistance. Highly penicillin resistant pneumococci are known to express low-affinity PBPs, an altered MurM protein, and possess a highly branched cell wall structure. It has therefore been hypothesized that MurM, and thus branched muropeptides, are essential for resistance because they are better substrates for low-affinity PBPs. In this study, we found that neither the version of *murM* nor elevated levels of cell wall branching affected the resistance level. To further support this, we investigated whether branched muropeptide substrates compete better than linear versions with penicillin at the active site of low-affinity PBPs and quantified changes to the stem peptide composition of the resistant Pen6 strain in response to subinhibitory concentrations of penicillin. We found that the level of cell wall branching decreased during penicillin exposure. Together our results do not support the idea that elevated levels of branched muropeptides (more active MurM) are important for either the function of low-affinity PBPs or the cell’s response to penicillin. Nevertheless, since a functional MurM enzyme is important for resistance, we speculate that it might indirectly influence other functions related to cell wall synthesis and remodelling needed for a resistant phenotype.

**Importance:** A fundamental understanding of the mechanisms behind antibiotic resistance is needed to find strategies to extend the clinical relevance of existing drugs. This study explores the relationship between cell wall composition and penicillin resistance in *Streptococcus pneumoniae*. Here we confirm that branched peptide crosslinks in the cell wall are crucial for resistance but found no correlation between elevated branching levels and resistance. Our data suggest that the function of low-affinity penicillin binding proteins is not influenced by the lack of branched cell wall precursors. Instead, a branched cell wall might contribute to resistance via other cell wall biosynthesis and remodelling mechanisms. These insights could offer new perspectives on why a branched cell wall is important for penicillin resistance in pneumococci.

## Introduction

*Streptococcus pneumoniae* (the pneumococcus) is an opportunistic human pathogen that colonizes the upper respiratory tract and can cause a wide range of infections including sinusitis, otitis media, community-acquired pneumonia, bacteremia and meningitis (Henriques-Normark & Tuomanen, 2013; Kadioglu et al., 2008). Respiratory tract infections are commonly treated with penicillins (β-lactam antibiotics), which inactivate the transpeptidation activity of penicillin binding proteins (PBPs) (Narciso et al., 2024; Tipper & Strominger, 1965; Zapun et al., 2008). PBPs are enzymes that contribute to synthesizing the peptidoglycan layer surrounding the cytoplasmic membrane by polymerising disaccharides of *N*-acetyl-glucosamine (GlcNAc) and *N*-acetyl-muramic acid (MurNAc) via transglycosylase reactions (Vollmer et al., 2008). Pentapeptides (L-Ala-D-iGln-L-Lys-D-Ala-D-Ala) attached to MurNAc are used by PBPs to incorporate the new glycan strands into the existing peptidoglycan layer by D,D-transpeptidation. PBPs recognize the D-Ala-D-Ala motif of the pentapeptides and link the carboxyl of D-Ala in fourth position of one glycan strand to the ε amino group L-Lys (or to L-Ala of an interpeptide bridge attached to L-Lys, see below) of the pentapeptide of an adjacent glycan strand (Sauvage et al., 2008; Vollmer et al., 2019). Since the β-lactams structurally mimics the natural D-Ala-D-Ala substrate of PBPs, covalent binding of penicillin to the active site for transpeptidation results in a weak bacterial cell wall that is depleted of inter-peptide bridges, leading to growth inhibition and/or cell lysis (Zapun et al., 2008). Unfortunately, the success of penicillin in treating pneumococcal infections is challenged by increasing prevalence of infections caused by penicillin resistant and non-susceptible strains (Gergova et al., 2024; Li et al., 2022). For instance, in China, it has been reported that the rate of resistant isolates is as high as 32%, and the number of non-susceptible isolates reaches up to 75% (Fu et al., 2019). The resistance to penicillin is mainly achieved by mutations in the genes encoding PBPs, which reduce their affinity to penicillin without disrupting their enzymatic activity (Hakenbeck et al., 2012). Fragments of genes encoding so-called low-affinity PBPs are typically acquired from other resistant pneumococci and relatives (*Streptococcus oralis* and *Streptococcus mitis*) via natural transformation resulting in mosaic-like *pbp* genes (Dowson et al., 1993; Potgieter & Chalkley, 1995; Reichmann et al., 1997).

*S. pneumoniae* has six different PBPs, of which five have transpeptidation activity (Sauvage et al., 2008). They are divided into class A and class B PBPs. Class A PBPs, namely PBP1a, PBP1b and PBP2a, are autonomous bifunctional enzymes that perform both transglycosylation and transpeptidation (Sauvage et al., 2008). Class B PBPs, PBP2x and PBP2b, are essential monofunctional transpeptidases that work alongside their cognate shape, elongation, division, and sporulation (SEDS) family glycosyltransferases FtsW and RodA, respectively (Berg et al., 2013; Cho et al., 2016; Emami et al., 2017; Kell et al., 1993; Meeske et al., 2016; Peters et al., 2014; Sjodt et al., 2020; Taguchi et al., 2019). Pneumococcal peptidoglycan is synthesised by a combination of septal and peripheral synthesis, where FtsW/PBP2x is found in the divisome and RodA/PBP2b in the elongasome (Berg et al., 2013; Massidda et al., 2013; Tsui et al., 2014). The class B PBP/SEDS complexes are responsible for synthesising the primary peptidoglycan, whereas class A PBPs are believed to exist both in the divisome and elongasome being important for repair, maintenance, and maturation of the peptidoglycan layer (Straume et al., 2021; Vigouroux et al., 2020). Expression of low-affinity versions of PBP2x, PBP1a and PBP2b has been found to be the main cause of β-lactam resistance in pneumococci (Grebe & Hakenbeck, 1996; Kosowska et al., 2004; Nichol et al., 2002; Sanbongi et al., 2004; Stanhope et al., 2008).

Although low-affinity PBPs are a prerequisite for penicillin resistance development in pneumococci, several studies have shown that additional proteins or genetic factors influence the resistance level (Chesnel et al., 2005; Crisóstomo et al., 2006; Dias et al., 2009; Gibson et al., 2022; Grebe et al., 1997; Guenzi et al., 1994; Huang et al., 2018; Kobras et al., 2023; Sauerbier et al., 2012; Schweizer et al., 2017; Smith & Klugman, 2001; Soualhine et al., 2005; Todorova et al., 2015; Tran et al., 2011). One of the most famous examples of this is the MurM protein. MurM attaches an L-Ala or L-Ser residue to the ε amino group of L-Lys of lipid II. This is followed by addition of an invariant L-Ala by MurN, resulting in a so-called branched muropeptide (GlcNAc-MurNAc-peptide) having Ala/Ser-Ala attached to the L-Lys ε amino group (Filipe et al., 2000a; Lloyd et al., 2008). In 1990, Garcia-Bustos and Tomaz reported that highly resistant pneumococcal isolates exhibited a completely different composition of peptidoglycan, shifting from a structure mostly consisting of linear peptide crosslinks between the glycan chains, to a peptidoglycan layer composed mostly of branched crosslinks. The elevated levels of branched peptidoglycan was found to be caused by expression of a mutated version of MurM (86% identity with MurM of the sensitive laboratory strain R6) which have been shown to be more active compared to a sensitive wild-type version (Lloyd et al., 2008). Multiple studies have found that *murM* mutations are transferred as a response to penicillin selection pressure *in vitro* (Garcia-Bustos et al., 1988; Gibson et al., 2022; Sauerbier et al., 2012; Smith & Klugman, 2001). Deletion of *murM* in resistant strains re-sensitizes them to penicillin, and this gene is therefore described as a resistance determinant in highly resistant isolates (Filipe & Tomasz, 2000b; Sauerbier et al., 2012; Weber et al., 2000). It has been proposed that low-affinity PBPs have increased preference for the branched lipid II precursor, which somehow hinders binding of β-lactams to the transpeptidation site (Garcia-Bustos & Tomasz, 1990). However, the correlation between MurM version, the level of branched muropeptides and a resistant phenotype is not fully understood. For instance, several studies have found that many resistant and non-susceptible strains express a normal *murM* version (Cafini et al., 2005; Chesnel et al., 2005; Davies et al., 2012; del Campo et al., 2006; Filipe et al., 2000c; Soriano et al., 2008) and expression of a mutated *murM* has been found in a sensitive strain (Schweizer et al., 2017). Additionally, transformation of a different *murM* versions into strains expressing low-affinity PBPs did not influence resistance (Filipe et al., 2001a; Plessis et al., 2002). Consequently, the precise mechanism by which MurM contributes to resistance is insufficiently characterized.

In this study, we have investigated the effect of cell wall branching in relation to penicillin resistance in pneumococci. Our results confirmed that increased activity of MurM directly correlates with more branched stem peptides in the peptidoglycan, but elevated peptidoglycan branching did not seem to influence resistance to penicillin. We also investigated the possibility of low-affinity PBPs preferring branched muropeptides over linear as a mechanism to reach high levels of penicillin resistance, but we could not find evidence supporting this. Undoubtedly, MurM is essential for penicillin resistance in pneumococci, however, our data opens for new interpretations as to why MurM has this function. We speculate that loss of resistance in a Δ*murM* mutant might not be because the PBPs become less efficient enzymes, but rather because complete absence of branched muropeptides influence other processes in cell wall synthesis and remodelling that are required for a resistant phenotype.

## Results

### Engineering a Penicillin G resistant pneumococcus: mosaic *pbp* genes and *murM* prove insufficient to develop a highly resistant phenotype

To understand how different PBPs contribute to a resistant phenotype, the penicillin susceptible *S. pneumoniae* strain R6 (Penicillin G [PenG] MIC_50_ = 0.032 µg/mL), hereafter named Wt, was transformed sequentially with amplicons of *pbp2x*, *pbp1a*, and *pbp2b* derived from the penicillin resistant *S. oralis* Uo5 (PenG MIC_50_ = 4.4 µg/mL), followed by selection on PenG (**Figure 1A**). By this strategy, we obtained mutants carrying low-affinity versions of PBP2x, PBP1a and PBP2b that allowed us to study the influence of each low-affinity PBP on resistance level, growth and stem peptide composition. The PBPs were verified to have low affinity for penicillin by reduced binding to Bocillin FL (**Figure 1C**) and MIC_50_ values of the resulting *pbp* mutants (and other strains) are summarized in **Table 1**. We found that low-affinity PBPs were acquired in a particular order when selecting for transformants with decreased PenG susceptibility. We were only able to transfer *pbp2x*^Uo5^ into the Wt using this method, while repeated attempts to transfer only *pbp1a*^Uo5^ or *pbp2b*^Uo5^ alone yielded no transformants. Furthermore, simultaneous transformation with all three *pbp* amplicons (*pbp2x*^Uo5^, *pbp1a*^Uo5^ and *pbp2b*^Uo5^) gave only rise to transformants with a mosaic *pbp2x*. In agreement with previous research (Plessis et al., 2002; Smith et al., 1993), these findings indicate that *pbp2x* must be the first to mutate. One transformant (strain MH10) expressing a low-affinity version of PBP2x (mosaic version of *pbp2x* comprising sequences from both R6 and Uo5 hereafter called *pbp2x^mos^*) was further transformed with either *pbp1a*^Uo5^ or *pbp2b*^Uo5^. A *pbp2x^mos^pbp1a^mos^* mutant was easily obtained (MH56), while only a few transformants were obtained for the *pbp2x^mos^pbp2b^mos^* double mutant (MH68) despite multiple attempts. This indicates that mutations in *pbp1a* more readily resulted in decreased penicillin susceptibility. After this, *pbp2b*^mos^ was easily created in the *pbp2x^mos^pbp1a^mos^*double mutant giving rise to a triple mutant (MH83) harbouring low-affinity version of all three PBPs important for resistance (**Figure 1A-B**). Considering the observed transformation pattern, we suggest that pneumococci acquire low affinity PBPs in the following order in response to PenG selection pressure: first mutations in *pbp2x*, followed by *pbp1a* and lastly *pbp2b* (**Figure 1A**). The MIC_50_ value increased 8-fold from Wt (MIC_50_ = 0.032 µg/mL) to the *pbp2x^mos^pbp1a^mos^pbp2b^mos^*triple mutant (MIC_50_ = 0.50 µg/mL) (**Figure 2B**). A recent study by Gibson et al. (2022) showed that during amoxicillin selection, the likely order of low-affinity PBP incorporation is *pbp2x* followed by *pbp2b* and lastly *pbp1a*. Numerous studies have proposed various orders in which pneumococci acquire low-affinity version of different PBPs (Fani et al., 2014; Gibson et al., 2022; Plessis et al., 2002; Reichmann et al., 1996; Smith & Klugman, 1998; Smith et al., 1993). A commonality among these is that *pbp2x* is the first PBP acquired. This makes sense considering the fact that PBP2x is essential and has the highest affinity to β-lactams among the PBPs in pneumococci (Kocaoglu et al., 2015).

**Figure 1.**
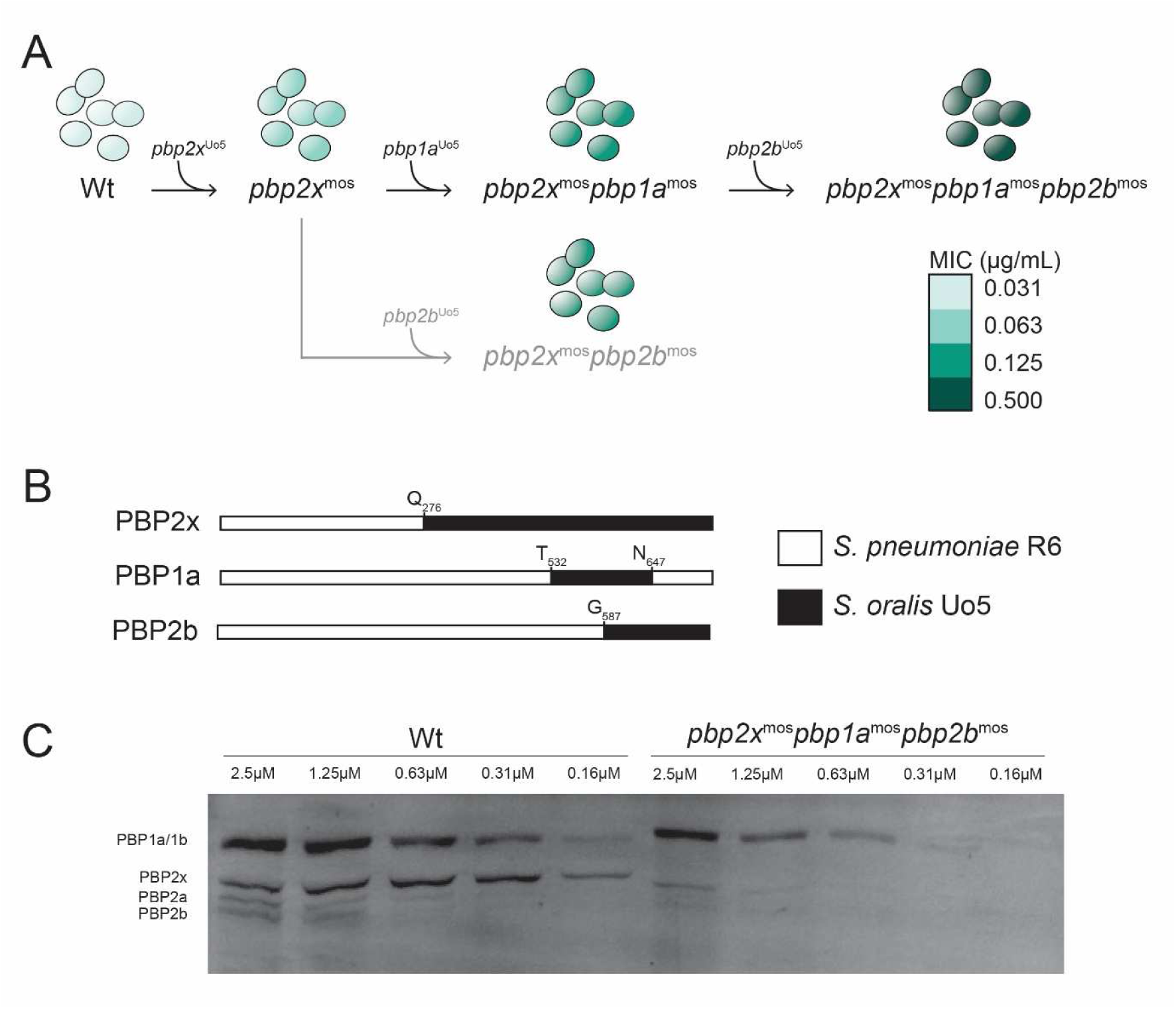
Transformation pattern and suggested order of mosaic *pbp* incorporation. (A) Wt was transformed with individual *pbp* genes from a highly resistant *S. oralis* strain (Uo5), resulting in mutants with various combinations of mosaic *pbps* (B). The incorporation of mosaic *pbps* increased the MIC_50_ value (the colours correspond to the mutants MIC_50_). MIC_50_ values (Table 1) represent the penicillin concentration that inhibited ≥ 50% of the maximum OD_550nm_. The suggested order of incorporated *pbps* is *pbp2x*, followed by *pbp1a* and finally *pbp2b*. (C) SDS-PAGE separated PBPs labelled with Bocillin FL confirmed that the mosaic PBPs had reduced affinity to penicillin. The concentrations of Bocillin FL used for labelling are indicated.

**Figure 2.**
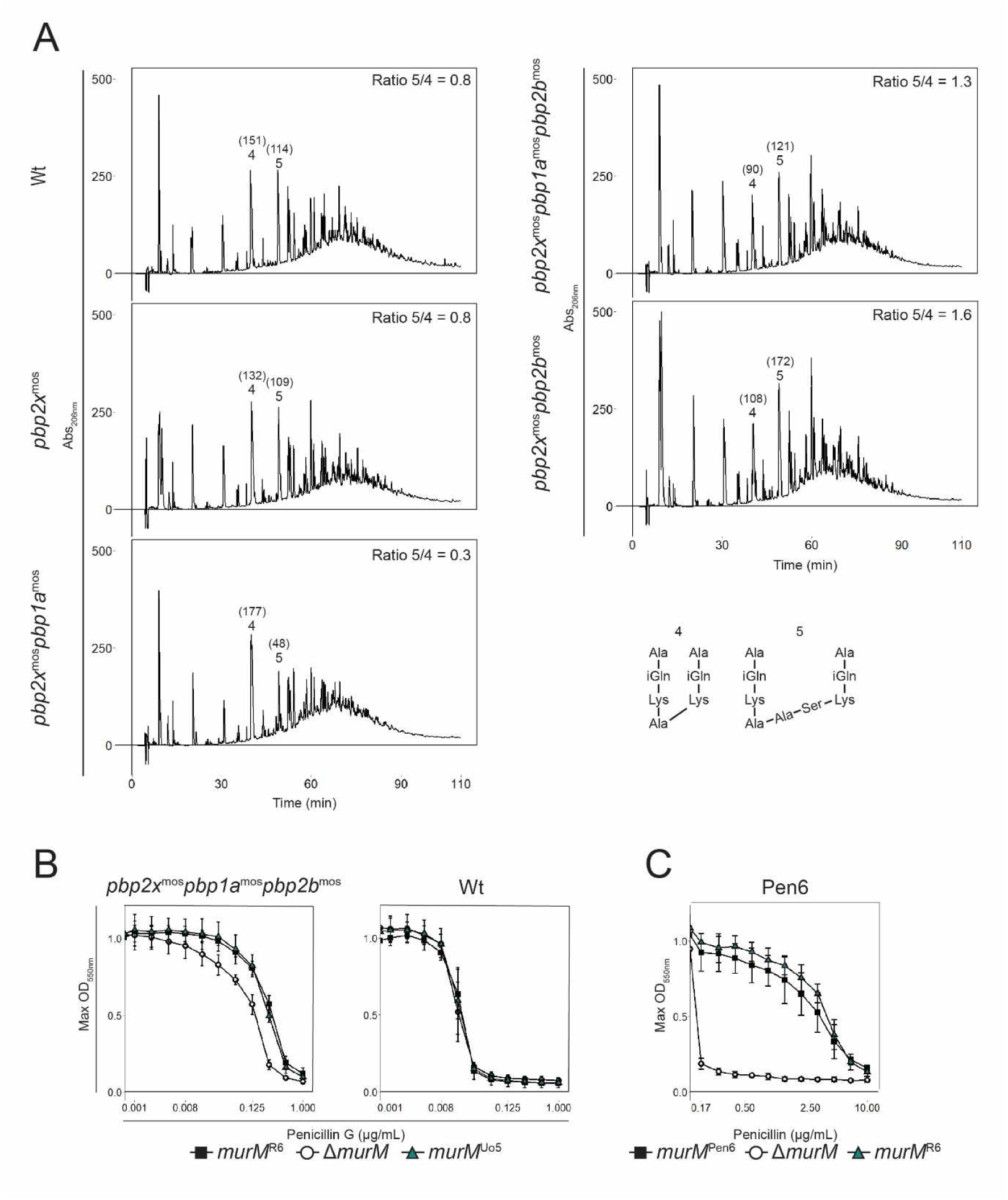
Stem peptide profiles of mutants carrying different low-affinity PBPs and PenG resistance profiles of different *murM* mutants. (A) Stem peptides of Wt and mutants expressing different combinations of low-affnity PBPs as indicated to the left of each graph. The area (mAU*min) of stem peptide 4 and 5, representing linear (Tetra-Tri) and branched (Tetra-Tri(SA) peptide crosslinks respectively, are shown in brackets and the peptide 5/4 ratios are indicated in the top right corners. Introduction of *pbp2b^mos^* shifted the proportion between branched and linear crosslinks. (B and C) The MIC curves of Wt, the *pbp2x^mos^pbp1a^mos^pbp2b^mos^*mutant and Pen6 display maximum OD_550nm_ obtained at each PenG concentration. For Wt and the triple mutant a two-fold dilution series of PenG starting at 1 µg/mL was used, and for Pen6 a 1.5-fold dilution series started at 10 µg/mL. Standard deviations were calculated from three biological replicates. Introducing *murM^Uo5^* to the triple mutant and the Wt had no effect on MIC. While deletion of *murM* dramatically reduced the MIC_50_ in Pen6, replacing it with the original version (*murM^R6^*) restored the high MIC_50_ of Pen6.

**Table 1.** MIC_50_ values of the different strains.

| Strains | <sup>a</sup>MIC<sub>50</sub> (μg/mL) | <sup>a</sup>MIC<sub>50</sub> (μg/mL)<br>(0.2 μM ComS) |
| --- | --- | --- |
| *S. oralis* Uo5 | 4.4 |  |
| RH425 (Wild-type) | 0.031 |  |
| Wt, $\Delta murM$ | 0.016 | |
| Wt, $murM^{Uo5}$ | 0.031 | |
| Wt, $pbp2x^{mos}$ | 0.063 | |
| Wt, $pbp2x^{mos}pbp1a^{mos}$ | 0.13 | |
| Wt, $pbp2x^{mos}pbp2b^{mos}$ | 0.13 | |
| Wt, $pbp2x^{mos}pbp1a^{mos}pbp2b^{mos}$ | 0.50 | |
| Wt, $pbp2x^{mos}pbp1a^{mos}pbp2b^{mos}, \Delta murM$ | 0.25 | |
| Wt, $pbp2x^{mos}pbp1a^{mos}pbp2b^{mos}, murM^{Uo5}$ | 0.25 | |
| Wt, $P_{comX}::murM^{R6}$ | 0.031 | 0.031 |
| Wt, $P_{comX}::murM^{Uo5}$ | 0.031 | 0.031 |
| Wt, $pbp2x^{mos}pbp1a^{mos}pbp2b^{mos}, P_{comX}::murM^{R6}$ | 0.50 | 0.50 |
| Wt, $pbp2x^{mos}pbp1a^{mos}pbp2b^{mos}, P_{comX}::murM^{Uo5}$ | 0.50 | 0.50 |
| Pen6 | 4.4 |  |
| Pen6, $\Delta murM$ | ≤0.17 | |
| Pen6, $murM^{R6}$ | 4.4 | |
| Pen6, $\Delta pbp1b$ | 4.4 | |
| Pen6 $\Delta pbp2a$ | 6.7 | |
| Pen6 $\Delta pbp1a$ | ≤0.17 | |
| Pen6 $\Delta eloR$ | ≤0.17 | |
| Pen6 $\Delta eloR\Delta pbp2b$ | 0.26 | |
| Pen6 $\Delta murM\Delta rsh$ | ≤0.17 | |
| Pen6 $P_{comX}::alaRS_{editing}, \Delta murM$ | ≤0.17 | ≤0.17 |

<sup>a</sup>The MIC<sub>50</sub> value was determined by the PenG concentration that inhibited ≥ 50% of the maximum OD<sub>550</sub>.

We also examined cell morphology, growth and stem peptide composition for each mutant harbouring low-affinity PBPs. Mutations in the PBP enzyme which decrease its affinity for penicillin, could possibly impact its ability to bind its natural substrate (i.e., lipid II and SEDS-produced glycan strands), potentially resulting in an impaired and/or altered cell wall synthesis. Such mutations have often been speculated to generate fitness disadvantages for the bacteria and several studies have observed such effects *in vitro* (Albarracın Orio et al., 2011; Barcus et al., 1995; Gibson et al., 2022) and *in vivo* (Trzciński et al., 2006). Our triple mutant did not display any defects with regard to cell size, morphology and growth (**Figure S1**). However, the *pbp2x^mos^pbp1a^mos^* double mutant displayed a small decrease in growth rate, which was rescued when *pbp2b^mos^* was introduced to this mutant. Interestingly, we observed that when a mutant was carrying *pbp2b^mos^*, the ratio of branched to linear dimer crosslinks were shifted (**Figure 2A**), indicating that expression of low-affinity PBP2b leads to an increase in branched peptide cross-links in the cell wall. This phenomenon is further elaborated in a later section.

Although the *pbp2x^mos^pbp1a^mos^pbp2b^mos^*triple mutant expressed low-affinity versions of all PBPs important for resistance, the MIC_50_ value (0.50 µg/mL) remained relatively low compared to the donor *S. oralis* Uo5 (4.4 µg/mL). This is consistent with previous studies, which found that transfer of additional genomic DNA was needed in addition to *pbp*s to reach the donor level of resistance (Fani et al., 2014; Plessis et al., 2002; Smith & Klugman, 2000; Smith & Klugman, 2001; Todorova et al., 2015). Several studies have proposed that a mutated version of the *murM* gene is required to reach a high MIC value (Gibson et al., 2022; Sauerbier et al., 2012; Smith & Klugman, 2001). We hypothesized that introduction of the altered MurM of *S. oralis* Uo5 (50% identity with R6 MurM) would further elevate the resistance level of our triple mutant. To our surprise, replacing *murM^R6^* with *murM^Uo5^* (strain RSG75) did not change the MIC value (**Figure 2B** and **Table 1**). Also, introducing *murM^Uo5^* to the Wt (RSG73) did not influence the MIC_50_ value consistent with previous studies (*Filipe* et al., 2001b; Filipe & Tomasz, 2000b). Overall, these results suggest that the acquisition of low-affinity PBPs was not sufficient to gain a highly PenG resistant phenotype, and that the introduction of a mutated *murM* version did not further enhance the resistance level.

### Elevated expression of *murM* increases cell wall branching but not penicillin resistance

Since we observed that expression of *murM^Uo5^* in the Wt and *pbp2x^mos^pbp1a^mos^pbp2b^mos^* triple mutant did not increase the MIC_50_ value, we were curious to explore the presumed link between expression of a mutated version of *murM* and enhanced penicillin resistance. It has been speculated that the genetic elements encoding penicillin resistance are separable from the genetic elements responsible for the abnormal cell wall found in resistant isolates (genetic elements that later was found to be *murM*) (Severin et al., 1996). Further, supporting this, Filipe et al. (2001a) have previously shown that replacing the β-lactam selected mutated *murM* version in strain Pen6 with *murM^R36A^*(derived from a penicillin susceptible laboratory strain) had no effect on penicillin susceptibility, suggesting that the version of *murM* was unrelated to the resistance level. Despite these findings, this research has largely gone unnoticed in the research field. We repeated this experiment by replacement of *murM^Pen6^* with *murM^R6^* in Pen6 (RSG186). Corroborating previous data (Filipe et al., 2000a), the replacement mutant showed a great decrease in branched muropeptides in the cell wall (**Figure S2**), resembling that of our triple mutant (**Figure 2A**), but no change in MIC_50_ (**Figure 2C**). See **Figure S3** for overview of stem peptide structures. This supported the hypothesis that mutations in *murM* is not important for resistance itself, even though such mutations are selected for based on decreased penicillin susceptibility. It is, however, important to emphasize that expression of a MurM protein is important for penicillin resistance since the MIC_50_ value dropped ≥ 26-fold in a Δ*murM* mutant (**Figure 2C** and first described by Filipe & Tomasz (2000b)), but it seems that elevated levels of cell wall branching could be independent of a resistant phenotype.

To further explore a possible correlation between incorporation of branched muropeptides in the peptidoglycan and resistance levels we performed *murM* overexpression experiments. By placing *murM^R6^* behind an inducible promoter, it was overexpressed in Wt and the *pbp2x^mos^pbp1a^mos^pbp2b^mos^*triple mutant (RSG200 and MH144). We saw increased levels of branched muropeptides in the cell wall of both strains when *murM^R6^* was overexpressed (**Figure 3A** and **S4A**). The stem peptide composition exhibited increased similarity to that of Pen6 (**Figure S2**) as well as other highly resistant clinical strains (Garcia-Bustos & Tomasz, 1990). However, introducing a highly branched structured cell wall by *murM* overexpression did not influence the PenG MIC_50_ (**Figure 3B** and **S4C**). We also tested overexpression of *murM^Uo5^* (strain MH105 and MH138) and obtained similar effects (**Figure S4**). Taken together, these results strongly indicate that there is no direct correlation between increased levels of branched peptidoglycan and the resistance level, regardless of low-affinity PBPs or the variants of the MurM protein expressed.

**Figure 3.**
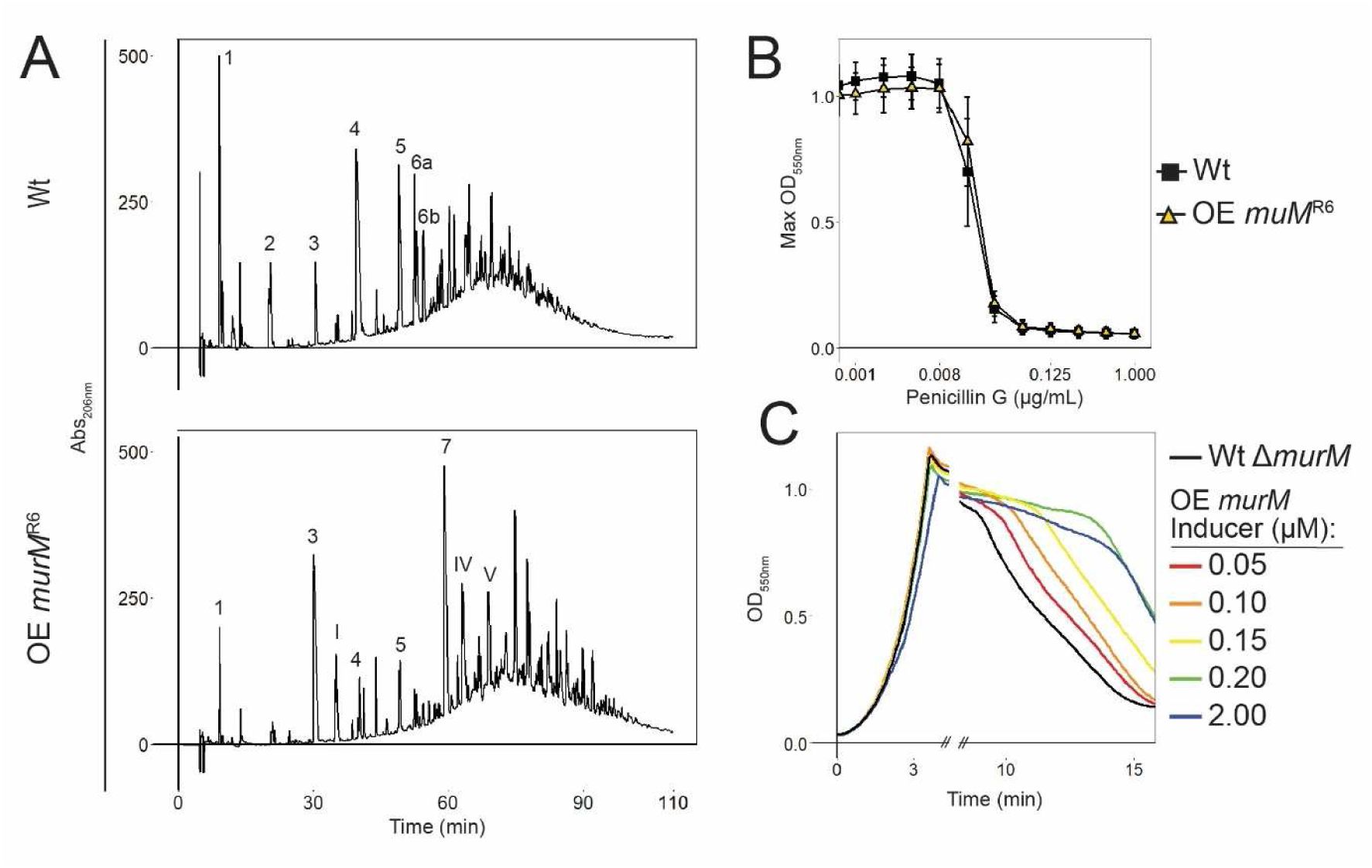
Effects of *murM* expression on the stem peptide profile, MIC and autolysis. (A) Stem peptides of Wt compared to Wt overexpressing (OE) *murM^R6^*. The stem peptides of the numbered peaks are illustrated in Figure S3. Overexpression of *murM* gave a shift from a less branched, to a highly branched cell wall structure. (B) The MIC curves display maximum OD_550nm_ at each PenG concentration of Wt compared to Wt overexpressing *murM^R6^*. Overexpression of *murM* had no effect on MIC_50_. Standard deviation was calculated from three biological replicates. (C) Autolysis was delayed in a dose dependent response to increasing expression levels of *murM^R6^*(concentrations of ComS inducer are indicated). The autolysis experiment was repeated tree times with similar results.

Of note, we observed that autolysis was delayed in cells overexpressing *murM* (**Figure S4C, D** and **S5**). Knockout of *murM* has been shown to cause earlier autolysis and increased sensitivity to the lytic effect of cell wall inhibitors (Filipe et al., 2000a; Filipe et al., 2002). We explored this effect further by making a Wt Δ*murM* mutant with inducible *murM^R6^* expression (RSG234) and found that reintroduction of *murM* expression delayed autolysis in a titratable manner (**Figure 3C**). These data suggested that an increased expression of MurM, that leads to a more branched cell wall structure, results in pneumococci with delayed autolysis.

### Overexpression of *murM* imposes toxic effects in Δ*murN* cells

Previous attempts to replace *murM* in strain R6 with *murM^Uo5^*did not succeed, and it has been suggested that *murM^Uo5^* is not tolerated in *S. pneumoniae* in the absence of low-affinity PBPs (Todorova et al. (2015). However, when we overexpressed this gene in our Wt strain, no growth defects were observed. To exclude that these cells were protected from the possible toxic effect of MurM^Uo5^ because they also expressed the native MurM^R6^, we repeated the experiment in a Δ*murMN* mutant (MH136). Surprisingly, we observed that the expression of *murM^Uo5^* had detrimental effects on these cells, resulting in severe growth defects with significantly slower growth and large morphological abnormalities (**Figure 4A**). Overexpression of *murM^Uo5^*was also toxic in a Δ*murMN* background of the *pbp2x^mos^pbp1a^mos^pbp2b^mos^*triple mutant (MH141, **Figure S6A**), showing that low-affinity PBPs did not alleviate the toxic effect of MurM^Uo5^ in the R6 strain. To test whether the toxic effect was unique to *murM^Uo5^*, we also overexpressed *murM^R6^*in Δ*murMN* backgrounds (MH146 and MH147) resulting in similar detrimental effects on growth (**Figure S6B**). Since *murM* overexpression was tolerated in Wt cells, we hypothesised that the absence of *murN* could be the determining factor for this observed toxicity. Indeed, in a Δ*murN* mutant, *murM* overexpression was shown to be toxic (MH149, **Figure 4B**). It has been shown that deletion of *murN* gives rise to muropeptides harbouring only a single amino acid (either serine or alanine) linked to the ε amino group L-Lys of the pentapeptide, instead of two (Filipe et al., 2000a). If *murM* is overexpressed in the absence of *murN*, it could lead to production of glycan chains containing high numbers of single amino acid branched stem peptides that might not be efficiently integrated into the cell wall, possibly explaining the observed growth defects. Analysis of stem peptide composition of Δ*murN* cells overexpressing *murM^Uo5^* showed significant changes (**Figure 4C**). This underlines the need for correctly structured peptidoglycan precursors to form a strong and resilient cell wall.

**Figure 4.**
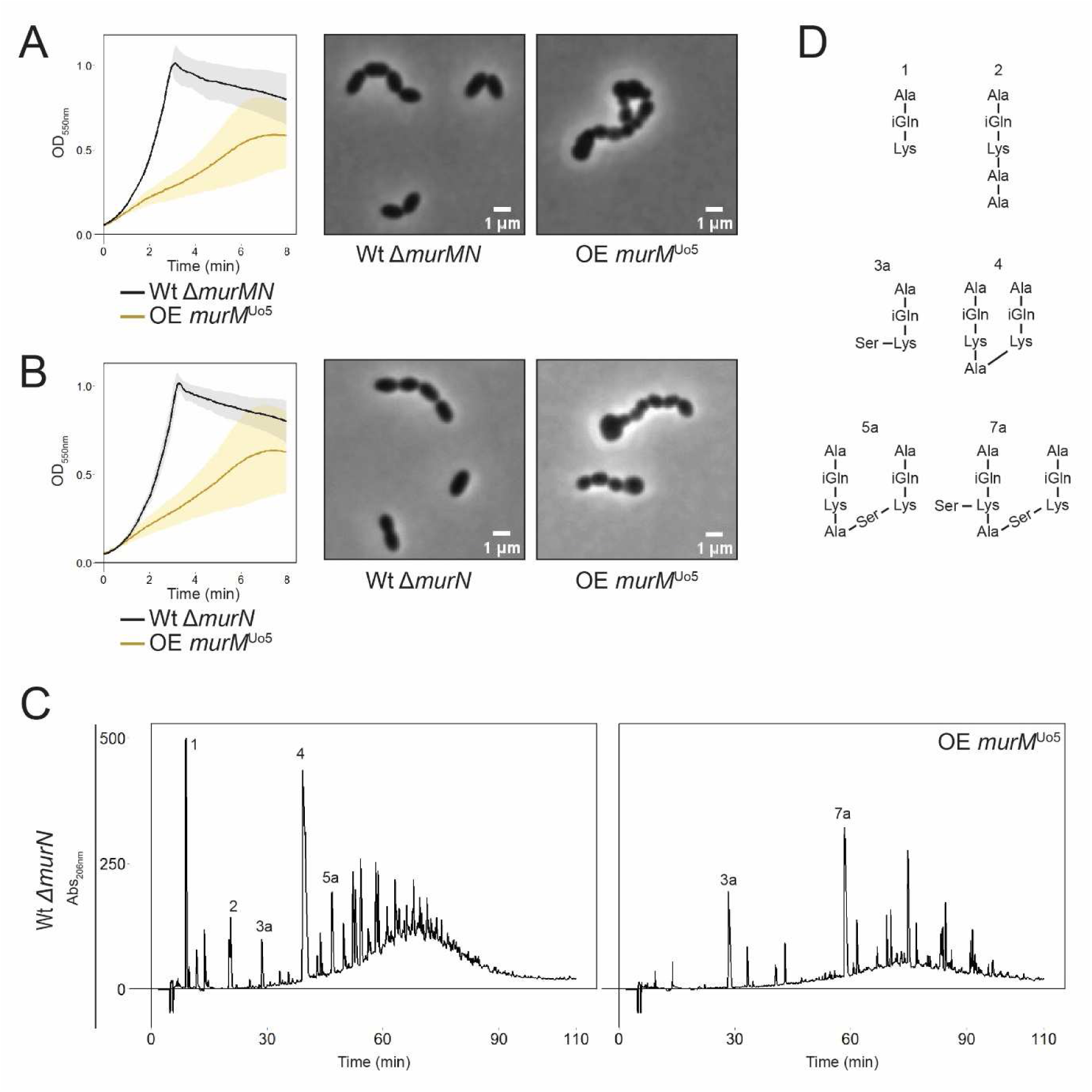
Impact of *murM^Uo5^*overexpression in Δ*murN* mutants. Overexpression (OE) of MurM^Uo5^ using 0.2 µM ComS inducer was not tolerated in MurN deficient cells (A, Δ*murMN* and B, Δ*murN*). Standard deviation was calculated from three biological replicates. Phase contrast microscopy of cells overexpressing *murM^Uo5^*at OD_550_ = 0.4 showed abnormal cell morphology when *murN* was deleted. (C) Overexpression of *murM^Uo5^* in the Δ*murN* mutant resulted in drastic changes to the cell wall stem peptide composition. (D) Structural representation of the stem peptides indicated in the chromatograms.

### Low-affinity PBPs’ impact on shaping a branched cell wall structure

We observed that mutants carrying a low-affinity version of PBP2b displayed increased levels of branched relative to linear stem peptide crosslinks (**Figure 2A**). Interestingly, a previous study also observed this shift in peptidoglycan composition in a Wt mutant depleted of *pbp2b* (Berg et al., 2013), indicating that a low-affinity PBP2b might be less functional. This is consistent with a study by Calvez et al. (2017) that demonstrated reduced transpeptidation activity of low-affinity PBP2b. These observations indicate that there could be a connection between the PBP2b enzyme functionality and incorporation of branched muropeptides in the cell wall. This phenomenon was explored by removing the low-affinity PBP2b from the Pen6 strain already having a high proportion of branched muropeptides in the cell wall. A double Δ*eloR*Δ*pbp2b* mutant (JM12) was used since *pbp2b* can be deleted in a Δ*eloR* background (Stamsås et al., 2017). No changes in the stem peptide composition were found in this mutant compared to Pen6 (**Figure 5A, S7A and Table S1**), indicating that low-affinity PBP2b and a Δ*pbp2b* mutant have the same effect on the cell wall.

**Figure 5.**
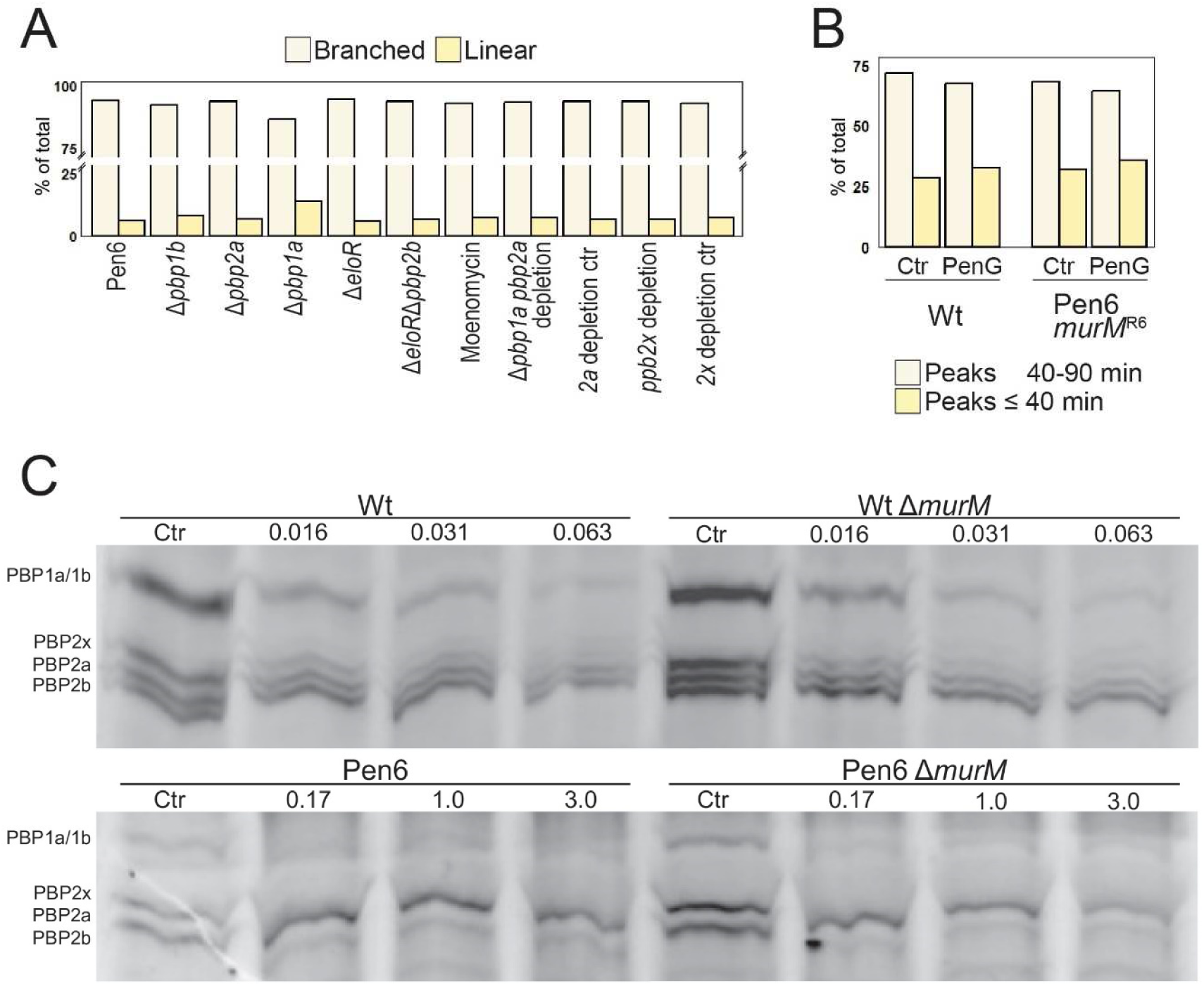
Effects of PBPs and PenG on stem peptide composition and PBP labelling. (A) Comparison of the relative levels (in percent) of branched and linear stem peptides in Pen6 cells in which PBPs are inactivated. The diagram is made from values presented in Table S1. (B) Total peak areas of linear (≤ 40 min) and branched stem peptides (40-90 minutes) for Wt and Pen6 *murM^R6^* treated with subinhibitory concentrations of PenG (See Figure S8 for chromatograms). Wt was treated with 0.016 µg/mL and Pen6 *murM^R6^*with 3 µg/mL PenG. Non-treated cells served as control (Ctr). Peak areas eluted after 40 minutes represent mostly branched stem peptides (although directly linked trimers are eluted here, they represent a minor fraction of the R6 cell wall strain (Bui et al., 2012)). The branched non-crosslinked stem peptides (peak 3 and I) eluting before 40 minutes were included in the percent of total branched. The level of branched stem peptides decreased relative to linear after exposure to sublethal concentration in both strains. (C) Bocillin FL labelling of PBPs in Wt and Pen6 and their respective Δ*murM* mutants after treatment of the cells with different concentrations of PenG (µg/mL indicated). Untreated cells prior to labelling were used as control (Ctr). Exponentially growing cultures (OD_550nm_ = 0.2) were treated with PenG for 30 minutes, excess antibiotic was removed by washing the cells prior to cell lysis and labelling with a final concentration of 15 µM Bocillin FL. The PBPs were separated using SDS-PAGE and visualised by fluorescence imaging (524/572 nm). No differences in fluorescence intensity of the PBPs were observed for the Δ*murM* mutants when compared to their respective parental strain.

Considering that other PBPs might compensate for PBP2b by incorporating more branched muropeptides, we next questioned whether all pneumococcal transpeptidases crosslink both linear and branched muropeptides with equal efficiency, or if they possess distinct selectivity towards these substrates. We explored this through various experiments, each involving inhibition or removal of one or more PBPs from the Pen6 strain followed by stem peptide analyses. The results from these analyses are summarized in **Figure 5A** and **Table S1** and are based on stem peptide profiles shown in **Figure S7**. The class A PBPs (PBP1a, PBP1b and PBP2a) were inhibited by addition of Moenomycin, an antibiotic specifically blocking these enzymes by inhibiting their transglycosylase activity. In addition, single Δ*pbp1a*, Δ*pbp1b* and Δ*pbp2a* knockouts and a double Δ*pbp1a*, *pbp2a* depletion mutant (since it has been proposed that these might have overlapping functions (Hoskins et al., 1999; Paik et al., 1999)) was analysed (strains JM9, JM5, JM6 and RSG208, respectively). Stem peptides of PBP2x depleted Pen6 was examined by using a *pbp2x* depletion mutant (AW594). However, none of these mutations or treatments resulted in any major changes to the stem peptide composition suggesting that neither PBP is solely responsible for incorporation of branched muropeptides. The lack of changes in the stem peptide composition could also indicate that the PBPs can compensate for each other, thereby masking any potential differences among stem peptide selectivity.

Although we did not see any major effects on cell wall stem peptide composition when deleting, depleting or inhibiting the different PBPs, we noticed that the Pen6 Δ*pbp1a* and Δ*eloR*Δ*pbp2b* mutant both displayed a substantial decrease in MIC_50_ (from 4.4 to ≤ 0.17 and 0.26 µg/mL, respectively). This confirms existing publications emphasizing that retaining the functionality of these PBPs, together with PBP2x, is critical for penicillin resistance (Grebe & Hakenbeck, 1996; Kosowska et al., 2004; Nichol et al., 2002; Sanbongi et al., 2004; Stanhope et al., 2008). It also shows that the expression of low-affinity PBP2b is important for resistance even if it might have reduced activity. It is important to mention that also the single Δ*eloR* mutant (RSG189) had the same MIC_50_ value as the Δ*eloR*Δ*pbp2b* mutant (JM12), implying that the effect is not solely due to *pbp2b*. Since EloR has been shown to be important for elongation of the cells and make *pbp2b* redundant (Stamsås et al., 2017), the effect of *eloR* deletion could be linked to absence of a functional PBP2b in the elongasome. Surprisingly, we found that Δ*pbp2a* caused an increase in MIC_50_ (from 4.4 to 6.7 µg/mL). Mutations in *pbp2a* have previously been observed in a few resistant strains (Chesnel et al., 2005; Sanbongi et al., 2004; Smith et al., 2005). Additionally, some studies have identified correlations between mutations in *pbp2a* and increased β-lactam resistance (Fani et al., 2014; Hakenbeck et al., 1998; Reichmann et al., 1996; Smith et al., 2005). It is possible that these mutations may not impact the enzyme’s affinity for penicillin, but rather its functionality, leading to lower susceptibility. As expected, no difference in resistance were seen when *pbp1b* was removed.

### The interplay between penicillin exposure and cell wall composition

Our results demonstrated that *murM* overexpression delayed autolysis (**Figure 3C, S4** and **S5**). Additionally, Δ*murM* mutants have been shown to cause earlier autolysis and increased sensitivity to the lytic effect of cell wall inhibitors (Filipe et al., 2000a; Filipe et al., 2002). This indicate that the presence of branched muropeptides in the cell wall is important for the cell wall integrity and susceptibility to cell wall hydrolases. We hypothesised that the protection from penicillin could result from low-affinity PBPs synthesizing a cell wall more resilient to murein hydrolases, i.e. increased levels of branched crosslinks upon exposure to penicillin. However, analyses of the stem peptide composition of the Wt and a Pen6 *murM^R6^* (RS186) strain (representing a sensitive and a resistant strain with Wt-like stem peptide composition) showed that subinhibitory concentrations of PenG reduced the amounts of branched crosslinks by 7.4 and 26.1%, respectively (**Table 2** and **Figure S8**). In comparison, linear crosslinked dimers (TriTetra) were reduced by 1.8% for both strains. We also compared the amounts of stem peptides eluting before ∼40 minutes representing linear peptides (except peptide 3 and I) with peptides eluting after 40 minutes (mostly comprising branched), which showed an increase of linear peptides (4.2% in Wt and 3.9% in Pen6 *murM^R6^*) and a corresponding reduction in branched upon PenG exposure (Figure 5B). Rejecting our hypothesis, these results indicate that the PBPs collectively incorporate less branched muropeptides into the cell wall during penicillin exposure.

**Table 2.** Stem peptide composition of Wt and a Pen6 *murM^R6^* mutant grown with subinhibitory concentrations of PenG.

| Peak | Peptide characteristics | Wt |  | Pen6 *murM<sup>R6</sup>* |  |
| --- | --- | --- | --- | --- | --- |
|  |  | Ctr | <sup>a</sup>0.016µg/mL | Ctr | <sup>a</sup>3.0 µg/mL |
| 1 | Linear Monomer | 16.8 | 12.6 | 19.9 | 4.8 |
| 2 | Linear Monomer | 3.8 | 16.5 | 3.0 | 40.3 |
| 3 | Branched Monomer | 6.9 | 4.5 | 9.4 | 2.0 |
| I | Branched Monomer | 1.5 | 1.1 | 2.2 | 0.4 |
| 4 | Linear Dimer | 20.6 | 18.8 | 14.0 | 12.2 |
| II | Branched Monomer | 0.9 | 3.7 | 1.1 | 12.2 |
| III | Branched Monomer | 0.4 | 1.1 | 0.3 | 4.0 |
| 5 | Branched Dimer | 15.8 | 15.3 | 18.4 | 9.6 |
| 6a | Branched Dimer | 7.3 | 5.8 | 6.3 | 2.1 |
| 6b | Branched Dimer | 5.8 | 4.2 | 3.9 | 0.6 |
| 7 | Branched Dimer | 3.9 | 2.0 | 5.2 | 0.8 |
| IV | Branched Dimer | 4.3 | 3.9 | 3.7 | 3.2 |
| 8 | Branched Trimer | 5.2 | 3.7 | 3.7 | 1.5 |
| 9 | Branched Trimer | 1.0 | 2.0 | 1.0 | 4.7 |
| V | Branched Dimer | 4.2 | 3.9 | 5.9 | 0.8 |
| VI | Branched Dimer | 1.4 | 0.7 | 1.6 | 0.4 |
| VII | Branched Trimer | 0.2 | 0.1 | 0.2 | 0.2 |
| VIII | Branched Trimer | 0 | 0 | 0.3 | 0.2 |
| IX | Branched Trimer | 0 | 0 | 0 | 0 |
| **Total (%)** |  | 100 | 100 | 100 | 100 |
| **Monomers (%)** |  | 30.4 | 39.4 | 35.8 | 63.7 |
| **Oligomers (%)** |  | 69.6 | 60.6 | 64.2 | 36.3 |
| **Linear peptides (%)** |  | 41.2 | 47.9 | 36.9 | 57.2 |
| **Branched peptides (%)** |  | 58.8 | 52.1 | 63.1 | 42.8 |
| **B/L peptides** |  | 1.4 | 1.1 | 1.7 | 0.7 |
| **Linear crosslinked peptides (%)** |  | 20.6 | 18.8 | 14.0 | 12.2 |
| **Branched crosslinked peptides (%)** |  | 49.1 | 41.7 | 50.2 | 24.1 |
| **B/L crosslinked peptides** |  | 2.4 | 2.2 | 3.6 | 2.0 |

<sup>a</sup>Final concentrations of PenG used to treat Wt and Pen6 *murM<sup>R6</sup>*.

While the overall level of branched stem peptides decreased, some peptides (II, III, 9) showed increased amounts for both the Wt and the Pen6 *murM^R6^* strain (**Table 2** and **Figure S8**), indicating that certain types of branched peptide bridges are more readily formed in the peptidoglycan during exposure to sublethal PenG concentrations. To further elaborate on this observation, we wondered whether the presence of branched muropeptide substrates could have a protective effect on the low-affinity PBPs’ susceptibility to penicillin. This could also explain why the presence of *murM* is highly important for penicillin resistance. To test this, a β-lactam competition assay using PenG and the fluorescently labelled β-lactam Bocillin FL were performed. Exponentially growing cells (Wt and Pen6 and their respective Δ*murM* versions, RH425 and RSG186) were pretreated with various PenG concentrations prior to cell lysis and Bocillin FL labelling. This enabled PenG to compete with the PBPs natural muropeptide substrates in actively growing cells. The PBPs not bound to PenG could subsequently be labelled with Bocillin FL. High degree of Bocillin FL labelling would indicate that the PBP was protected from PenG by its natural substrates. However, there were no clear differences in PBP-labelling between either Wt or Pen6 and their respective Δ*murM* mutants (**Figure 5C**) indicating that the absence of branched muropeptide precursors is not affecting the PBPs affinity to PenG.

### Could MurM’s role in the stringent response pathway promote protection from penicillin treatment?

A recent study showed that MurM can repress activation of the stringent response in *S. pneumoniae* (Aggarwal et al., 2021). This is a stress response, which in *S. pneumoniae*, has been shown to have a strong negative effect on cell growth, resembling stationary phase followed by LytA-mediated autolysis (Aggarwal et al., 2021; Hauryliuk et al., 2015). This is also the main outcome for pneumococci inhibited by β-lactams (Tomasz et al., 1970; Tomasz & Waks, 1975). Interestingly, MurM was found to prevent activation of the stringent response by removing serine from misaminoacylated Ser-tRNA^Ala^ (acidic stress induced) by adding it onto Lipid II precursors for incorporation into the peptidoglycan (illustrated in **Figure S9**). In this way the cell wall serves as a drainage system for removing toxic levels of Ser-tRNA^Ala^ molecules during stress. Deletion of one of the alarmone producers (RSH) rescued the Δ*murMN* phenotype (Aggarwal et al., 2021). This discovery prompted us to test if penicillin resistant strains are dependent on MurM due to its function to prevent accumulation of Ser-tRNA^Ala^ and activation of the stringent response, rather than for the changes to the cell wall composition. We tested this hypothesis by comparing PenG resistance of a Pen6 Δ*murM* mutant with (i) a double Δ*murM*Δ*rsh* mutant (RSG219) in which elevated Ser-tRNA^Ala^ levels will not activate the stringent response, and (ii) a Δ*murM* mutant overexpressing the editing domain of the AlaRS enzyme (RSG243), which removes serine from Ser-tRNA^Ala^. If resistance depends on MurM for moving serines of misaminoacylated Ser-tRNA^Ala^ into the peptidoglycan to prevent activation of the stringent response, we expected the mutants lacking *rsh* and the AlaRS overexpression mutants to display significantly higher resistance to PenG than the single Δ*murM* mutant. No differences in the MIC_50_ values were seen (**Figure S10**), indicating that reducing unfavourable levels of Ser-tRNA^Ala^ by incorporating Ser into the peptidoglycan via lipid II is not the main reason behind the essentiality of *murM* for penicillin resistance.

## Discussion

It is well established that the version of MurM expressed in pneumococcal isolates can influence the level of branched muropeptides that are incorporated into the peptidoglycan layer (Filipe et al., 2000a). Here, we demonstrated that this phenomenon can also occur by changing between PBPs with high and low affinity to β-lactams. More specifically, introduction of a low-affinity PBP2b increased the level of branched muropeptides in the cell wall even though the MurM was unaltered (**Figure 2A**). It suggested that low-affinity PBP2b either prefers branched muropeptides as substrate or that other PBPs incorporate more branched muropeptides to compensate for a possibly reduced transpeptidase activity of low-affinity PBP2b. Since deletion of low-affinity PBP2b in the highly resistant Pen6 strain (which expresses a low-affinity PBP2b and is known to have a high proportion of branched muropeptides in the cell wall), did not result in significant decrease of branched muropeptides (**Figure 5, S7A** and **Table S1**) it favoured a model where other PBPs incorporate more branched muropeptides to compensate for reduced transpeptidase activity of low-affinity PBP2b. This could also explain previous observations that depletion of the *pbp2b* gene resulted in increased branching (Berg et al., 2013). To our surprise, however, systematic inactivation or depletion of the other PBPs had minimal influence on the stem peptide composition. Thus, it seems that no single type of PBP is exclusively responsible for incorporating branched muropeptides into the cell wall of a resistant strain, but that it is rather a result of the combined action of several PBPs.

We observed that sequential transformations of genes encoding low-affinity PBPs from *S. oralis* Uo5 into *S. pneumoniae* R6 increased resistance to PenG, but it did not recreate the high-level resistance of the donor. We reasoned that the missing factor could be the highly mutated version of *murM* from *S. oralis* Uo5. However, replacement of *murM^R6^* with *murM^Uo5^* did not increase resistance (**Figure 2B**). This is in line with the results of Filipe et al. (2001a) (see also **Figure 2C** and **Table 1**), who did the opposite by reintroducing *murM* from a penicillin sensitive strain into Pen6 (a strain with a mutated *murM^Pen6^* acquired upon penicillin selection) without seeing a reduction in resistance, despite a decrease in the level of branched muropeptides. In fact, many studies show that both β-lactam resistant and non-susceptible clinical isolates express a normal *murM* version (Cafini et al., 2005; Chesnel et al., 2005; Davies et al., 2012; del Campo et al., 2006; Filipe et al., 2000c; Schweizer et al., 2017; Soriano et al., 2008), which demonstrate that high penicillin resistance is not dependent on a mutated *murM*. When analysing a selection of 81 resistant isolates, we found that 75% express a MurM with > 95% identity to wild type MurM (**Figure S11A**) also indicating that the correlation between resistance and *murM* mutations is not absolute. We also found that increasing the level of branched muropeptides in the cell wall by overexpressing *murM* did not influence the PenG MIC level in neither Wt nor the triple mutant (**Figure 3B**, **S4C-D**). This strengthens the notion that increased levels of branched muropeptides in the cell wall of pneumococci is not critical for the penicillin resistance function, and that the link between penicillin resistance and an altered version of MurM is more intricate than previously thought. Nevertheless, mutations in *murM* are selected for upon acquiring low-affinity PBPs in some strains. One could speculate that the benefit of acquiring a mutated *murM* might depend on the specific combination of low-affinity PBPs expressed or other features of the genetic background. Sequence analyses of PBPs and MurM have suggested a co-evolutionary link for a particular set of PBPs and MurM (Cafini et al., 2005; del Campo et al., 2006; Sauerbier et al., 2012). Our dataset was too limited for a formal analysis of MurM evolution at the level of individual GPSCs, but we did find that some GPSCs harboured multiple MurM alleles (**Figure S11B**).

Given our perception that the MurM version and increased amounts of cell wall branching is less important for the level of resistance, we tried to understand why Δ*murM* mutants of resistant strains become resensitized to penicillin. A commonly proposed explanation is that the low-affinity PBPs might have altered substrate specificity or require branched muropeptides for peptidoglycan synthesis. Since *murM* can be deleted in resistant strains with minimal effect on growth under normal laboratory conditions, this seems unlikely. Another hypothesis is that branched substrates can better compete with β-lactams for binding to the transpeptidation site. We tested this by investigating if the substrate specificity changed during penicillin exposure (**Figure S8** and **Table 2**) or if the presence of branched muropeptide precursors altered the penicillin affinity of the PBPs (**Figure 5C**). If branched muropeptides are better competitors to penicillin than the linear muropeptides, we would expect to see that the cell wall consists of more branched than linear peptide crosslinks during penicillin exposure. However, our results showed the opposite; that the cell wall of penicillin treated Pen6 *murM^R6^* contained less branched cross-links (24.1%) comparable to the non-treated control (50.2%, **Table 2**) indicating that such competition is not present. Since 3 µg/mL PenG was used, PBP1b and PBP2a are probably blocked, meaning that mainly PBP2x, PBP1a and PBP2b (low affinity versions) synthesized the peptidoglycan. This could indicate that low-affinity PBPs do not incorporate more branched crosslinks during penicillin exposure. The results from the Wt were comparable (41.7 % branched crosslinks in PenG treated culture and 49.1% in non-treated control), indicating that both high- and low-affinity PBPs incorporate mostly the same peptidoglycan precursors. Further supporting this result, our PenG-Bocillin FL competition assay did not show that branched muropeptide precursors were better competitors for the active site than linear muropeptides (**Figure 5C**). Together, the findings mentioned above indicate that branched precursors are not used to a larger extent during PenG treatment. In fact, the expression of *murM* has been reported to go down during penicillin exposure of the sensitive D39 strain (Tran et al., 2011), which is opposite of what would be expected if branched muropeptides becomes more important during penicillin treatment. Whether this is also the case for resistant strains are not known.

Since competition between β-lactams and muropeptides at the transpeptidation site might not be the reason for MurM’s essentiality for resistance, we investigated whether MurM is important due to its role in stress regulation, but did not find any evidence to support this (**Figure S10**). Also, to check whether it was the activity of MurM that is important for resistance or if it is the presence of the protein itself (e.g. as being a protein complex critical for resistance), we created catalytically inactive MurM mutants. Analysis of the 3D structure of MurM suggested residues K35, W38, R215 and Y219 to be essential for lipid II binding (York et al., 2021). By replacing *murM* in Pen6 with either *murM^K35A,W38A^*, *murM^R215A,Y219A^*, or a combination of all 4 mutations (strains AW627, AW628 and AW629) we saw that resistance dropped to that of a Δ*murM* mutant, indicating that MurM activity (not only presence) is essential for PenG resistance (**Figure S12**).

Although the present work as well as previous studies report that high level penicillin resistance is compatible with expression of a normal MurM and we show that increasing cell wall branching above the normal levels has little effect on resistance, *murM* is undoubtedly an important resistance determinant. Why some pneumococci continuously mutate *murM* both in penicillin-selective growth media and in their natural environment, despite that the effect appears to be minimal on resistance levels, remains elusive. Perhaps mutations in *murM* facilitate (or necessitate) the acceptance of specific *pbp* mutations, either spontaneous point mutations or acquired by natural transformation. There may also be other potential benefits to a mutated or more active MurM. For example, we demonstrated that overexpression of *murM* resulted in a more branched cell wall structure and delayed autolysis (**Figure 3A, S4** and **S5**). It is possible that branched muropeptides in the cell wall influences the cell’s ability to control autolysis by offering some protection from LytA or other cell wall hydrolases. A similar observation has been shown for the release of the major pneumococcal virulence factor Pneumolysin (Ply), which depends on the action of murein hydrolases (Greene et al., 2015). Release of Ply increased in Δ*murM* mutants, suggesting that linear cross-linked peptidoglycan is more susceptible for muralytic degradation. The idea that a branched cell wall is more resilient to uncontrolled degradation during β-lactam treatment has also been suggested by others, i.e. that an imbalance of activity between cell wall synthesising- and remodelling enzymes develops. In the absence of a branched peptidoglycan layer, such imbalance may lead to opening of the cell wall without sufficient material being inserted, potentially triggering further lysis by other peptidoglycan hydrolases (Calvez et al., 2017). The detrimental effects we observed for cells overexpressing *murM* in a Δ*murN* background (most probably resulting in increased levels of lipid II with only one amino acid branch) exemplifies how a mis-constructed peptidoglycan layer leads to critical cell stress (**Figure 4**).

In this work we set out to find new clues as to why *murM* is essential for β-lactam resistance in pneumococci. Our initial goal was to identify whether PBPs display higher preference for branched muropeptides in a resistant strain, but instead our data suggest a strong substrate promiscuity among the PBPs. Furthermore, we saw no direct correlation between increased cell wall branching and resistance levels or that branched muropeptides somehow protect low-affinity PBPs against penicillin. This suggest that the function of MurM for resistance could be linked to other cellular functions. Our results do not support MurM’s role as a stringent response regulator to be this function. Why MurM is essential for high penicillin resistance in pneumococci remains an open question. Binding kinetic studies of individual PBPs using penicillin in the presence of glycan strands composed of only linear or branched stem peptides could possibly confirm or reject the PenG-Bocillin FL competition assay performed here and give a definite answer if branched muropeptides are important for keeping penicillin out from the transpeptidase site. In addition, since the class B PBPs (PBP2x and PBP2b) depend on their cognate transglycosylases FtsW and RodA to obtain their substrates, it would be interesting to investigate how low-affinity PBPs work alongside their dedicated SEDS polymerases, which are not mutated in resistant strains. The length of the glycan chains has been shown to influence resistance to cell wall targeting antibiotics in *Staphylococcus aureus* (Chan et al., 2016; Komatsuzawa et al., 2002; Rebets et al., 2014). Whether the glycan chain length is significantly shorter in Δ*murM* mutants and if this is critical for resistance levels should be investigated in the future. The possibility that MurM could have additional roles e.g. influencing gene expression, protein activation or protein complex formation which is critical for a resistant phenotype should also be explored.

## Methods

### Bacterial strains and growth conditions

The strains used in this work are listed in **Table S2**. *S. pneumoniae* was grown at 37 °C in C-medium (Lacks & Hotchkiss, 1960) without shaking or on Bacto™ Todd Hewitt (TH) broth (Becton Dickinson) agar in an anaerobic chamber using Oxid AnaeroGenᵀᴹ (Thermo Scientific). When necessary, the following antibiotics were used: kanamycin (400 µg/mL), streptomycin (200 µg/mL) and PenG (concentrations indicated when appropriate). *S. oralis* was grown in Todd Hewitt broth at 37 °C without shaking.

### Construction of genetic mutants

Pneumococcal mutants were created by transformation with PCR products containing either a selection marker gene or a mutated version of a gene of interest flanked by ∼1000 bp regions up- and down-stream of the relevant position in the genome. Primers used to create transformation cassettes are listed in **Table S3**. In general, to make knockout mutants, the gene of interest was replaced by the Janus cassette (Sung et al., 2001). The Janus constructs were created by overlap extension PCR (Higuchi et al., 1988). Sequences corresponding to ∼1000 bp upstream and downstream of the gene to be deleted were amplified and fused with the 5’ and 3’-end of the Janus cassette, respectively. The same technique was used to create gene fusion or point mutations. To create mutants for ectopic gene expression (overexpression or gene depletion assays), the ComRS system for gene depletion (Berg et al., 2011) was used. To create gene depletion strains, P1::P*_comR_::comR* was placed between *amiF* and *treR* and the gene of interest was placed behind the P*_comX_* promoter located between *cspN* and *cspO*. Natural transformation of *S. pneumoniae* was used to introduce genetic changes. Exponentially growing cultures were diluted to OD_550nm_ of 0.05-0.1 and incubated at 37 °C for 15 minutes before a final concentration of 250 ng/mL competence-stimulating peptide 1 (CSP-1; NH_2_-EMRLSKFFRDFILQRKK-COOH) was added together with a final concentration of 100-200 ng/mL of transforming DNA. Following 120 minutes of incubation at 37 °C, 30 µL of the cell cultures were spread on TH agar plates containing the appropriate antibiotic and 0.2 µM ComS when required to drive expression of an essential gene. For selection on PenG, plates containing a 1.5x gradient of antibiotic just above and below the MIC_50_ concentration of the transformed strains were used. Knockout mutants were screened with PCR and introduction of mutations was confirmed with Sanger Sequencing.

### Gene overexpression- and depletion

Genes were ectopically expressed in various pneumococcal genetic backgrounds using the ComRS system (Berg et al., 2011). In brief, a gene of interest was placed behind P*_comX_* and gene expression was induced at OD_550nm_ = 0.05 by adding a final concentration of 0.2 µ M ComS inducer (NH_2_-LPYFAGCL-COOH). Concentrations other than 0.2 µ M are indicated. For gene depletion experiments, *pbp2a* or *pbp2x* was first placed behind P*_comX_* and the native gene was subsequently replaced by Janus. When the native gene was replaced, the growth medium was supplemented with 0.2 µM ComS. Cultures were grown in C-medium containing 0.2 µM ComS and exponentially growing cells (OD_550nm_ of 0.2-0.4) were harvested by centrifugation and washed three times with C-medium to remove excess ComS. The cells where diluted to an OD_550nm_ of 0.05 and a two-fold dilution series of the cells were made in a 96-well plate. Volumes of 100 µL diluted culture in the plate were further diluted by adding 200 µ L C-medium (depletion) or C-medium with ComS (control) to the wells. OD_550nm_ was measured every 5 minutes for 16-20 hours at 37°C using a Hidex Sense microplate reader. The growth measurements were used to calculate the inoculum required to obtain gene depletion effect at OD_550nm_ = 0.3-0.4 in 1 L cell cultures harvested for cell wall isolation.

### Growth experiments and microscopy

Exponentially growing cultures (OD_550nm_ = 0.2-0.4) were diluted to OD_550nm_ = 0.05 in C-medium (when relevant, ComS was added). Volumes of 300 µ L diluted cultures were transferred to a 96-well plate and growth were followed by measuring OD_550nm_ every 5 minutes for 16-20 hours using a Hidex Sense microplate reader. MIC_50_ assays were also performed in 96-well plates using a dilution series of PenG (sodium salt, Sigma Aldrich) in C-medium. Culture volumes of 260 µ L (OD_550nm_ = 0.05) were added to 40 µ L antibiotic dilution series in the plate. When relevant, the inducer peptide ComS was added. MIC_50_ was determined as the concentration of antibiotic that inhibited ≥50% of the average bacterial growth (maximum OD_550nm_ reached without PenG) using three biological replicates. For phase contrast microscopy the cultures were grown to OD_550nm_ = 0.4. The cells were examined using a Zeiss AxioObserver with an ORCA-Flash 4.0 V2 Digital CMOS camera (Hamamatsu Photonics) and a 100x phase-contrast objective. The microscope was operated using the ZEN Blue software. Images were prepared and analysed using the ImageJ software with the microbeJ plug-in (Ducret et al., 2016).

### Cell wall isolation and analysis

Bacteria were cultivated to OD_550nm_ = 0.4-0.5 in 0.5 or 1 L volumes of C-medium. When appropriate, antibiotics (concentrations indicated in the text) or inducer peptide (0.2 µ M ComS) was added to the culture at OD_550nm_ = 0.05. Cells were harvested at 10 000 *g* for 10 minutes. Cell wall was isolated and purified as previously described (Vollmer, 2007). The purified cell walls were digested with LytA as described previously (Straume et al., 2017). The stem peptides from 0.5 mg cell wall were separated by HPLC using a C_18_ reverse-phase column (Halo 160 Å ES-C18, 2.7 µm, 4.6 x 250 mm from Advanced Material Technology) coupled to a Dionex Ultimate 3000. Stem peptides were eluted using a 120-minutes linear gradient of acetonitrile from 0-15 % in 0.05% trifluoracetic acid. The flow rate was set to 0.5 mL/min. Peptides were detected at 206 nm. Eluted stem peptides were directly injected into a Velos Pro dual-cell 2D linear ion trap mass spectrometer for mass determination. Mass spectrometry was performed in positive mode using the XCalibur software. Mass spectrometry data was analysed in Freestyle software (Thermo Fisher Scientific).

### Bocillin-FL binding assay

Cultures of 10 mL were grown to an OD_550nm_ of 0.2 and harvested by centrifugation at 4000 *g* for 10 minutes. The cells were resuspended in 100 µ L sodium phosphate buffer (20 mM, pH = 7.2) with 0.2% Triton® X-100 (Sigma-Aldrich). The cells were lysed by adding purified LytA (25 µg/mL) following incubation at 37 °C for 10 minutes. The lysate was stored at −80 °C prior to analysis. Total protein concentration was estimated by measuring absorbance at 280 nm using NanoDrop 2000 (Thermo Scientific). For the PenG-Bocillin FL competition assay, the growing cultures were incubated with different concentrations of PenG at 37°C for 30 minutes prior to harvesting. The cells were then washed 3x with 1 mL PBS (pH 7.4) to remove unbound antibiotic. Labelling with fluorescently labelled penicillin (Bocillin FL) was performed by adding concentrations of Bocillin FL ranging from 15-0.16 µM to 15µ L of lysate (40 mg/mL) followed by incubation at 37 °C for 30 minutes. The labelled lysate was mixed 1:1 with 2x SDS sample buffer and 15 µ L of the sample was applied to SDS-PAGE. Electrophoresis was performed by applying 1.25 V/Cm^2^ for 15 minutes followed by 2.5 V/Cm^2^ until the migration front reached the end of the gel. Then electrophoresis was continued for another 90 minutes for sufficient separation of the labelled PBPs. The SDS-PAGE gel consisted of a 4 % stacking gel and 10 % separation gel previously described by Rutschman (2007). The labelled PBPs were visualised using Azure c400 imaging system (Azure Biosystems) at 524/572 nm and images were prepared using the ImageJ software.

### Analysis of MurM sequences

In order to assess MurM amino acid sequence variation in a larger panel of clinical *S. pneumoniae* isolates, a selection of genome assemblies were retrieved from the Global Pneumococcal Sequencing Project (GPS) database (https://www.pneumogen.net/gps/ - accessed 2024-06-14). We first downloaded assemblies of isolates with a reported penicillin MIC in the range of 4-16 mg/mL and which belonged to GPS clusters (GPSC) containing > 2 isolates exhibiting penicillin MICs in this range. If > 10 isolates were found for a single GPSC, 10 random isolates were selected. Next, we downloaded assemblies belonging to the same GPSCs, aiming to match the number of sequences, but with reported penicillin MICs < 0.064 mg/mL. However, we ended up with a smaller number of susceptible isolates, as we could not retrieve a matching number of low-MIC isolates for all GPSCs. Next, in order to retrieve MurM sequences, we matched the assemblies against the MurM amino acid sequence of D39 (representing a wildtype sequence) using blastx (Camacho et al., 2009), retrieving the best hit with a minimum alignment length = 300.

## Acknowledgements

This work was supported by grants from The Research Council of Norway (Grant No. 314720), the National Institute of Public Health and the Norwegian University of Life Sciences. The authors would like to acknowledge Zhian Salehian for technical assistance. We would also like to thank Thales de Freitas Costa for help with LC-MS analyses.

