## Supplementary material for "The effect of MurM and a branched cell wall structure on penicillin resistance in *Streptococcus pneumoniae*": Supplemetal material

### Supplementary figures

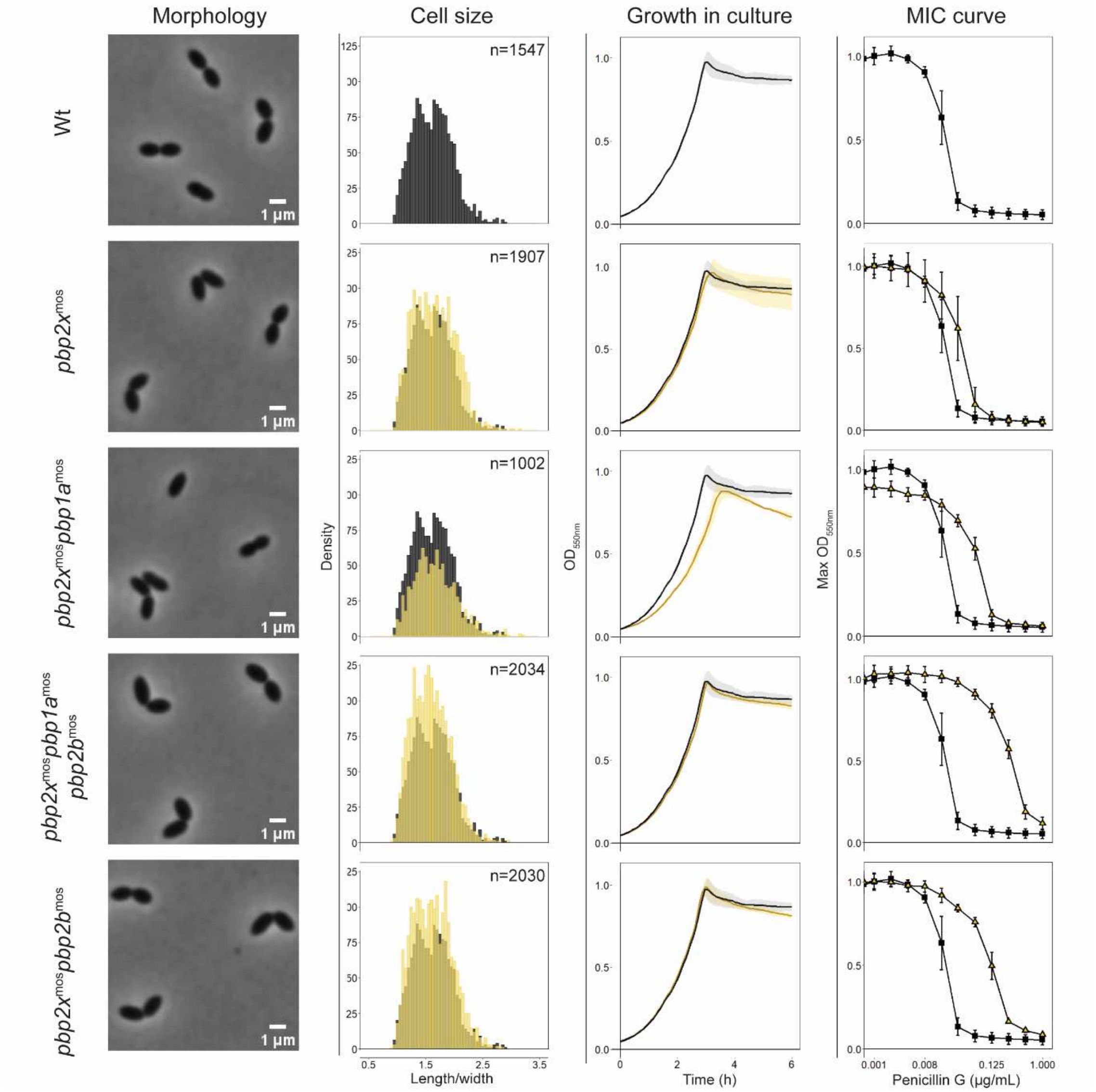


**Figure S1: Phenotypical characteristics of Wt and mutants with different combinations of low-affinity PBPs.** Cultures were diluted to an $OD_{550nm}$ of 0.05 and grown to an $OD_{550nm}$ of 0.4 for phase contrast microscopy. Images display a representative view of the different mutants. Cell size distribution was calculated based on length/witdh ratios, with meassurements performed by the ImageJ and MicrobeJ plug-in. The numbers of cells included in the analysis are indicated. Bacterial growth ($OD_{550nm}$) in liquid culture was measured continuously every 5 minutes for 16 hours (only the first 6 h are displayed). The MIC curves display maximum $OD_{550nm}$ at each PenG concentration (a two-fold dilution series starting at 1 µg/mL) with the $MIC_{50}$ value (**Table 1**) determined by the PenG concentration that inhibited ≥50% of the maximum $OD_{550nm}$. Wt measurements are shown in black, and the different low-affinity *pbp* mutants (specified to the left in the figure) are show in yellow. Standard deviation was calculated from three biological replicates.

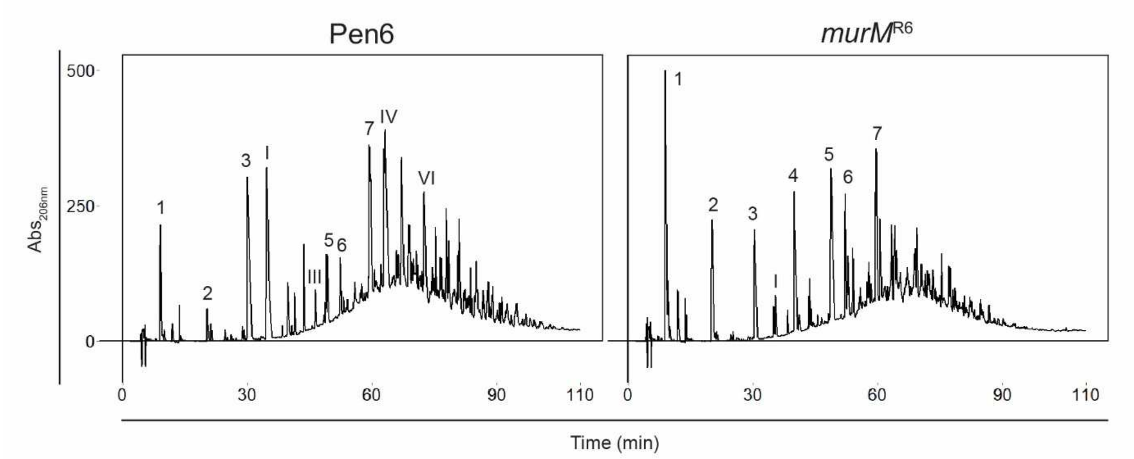


**Figure S2: Stem peptide analysis of peptidoglycan from Pen6 and a mutant in which the highly mutated *murM*$^{Pen6}$ has been replaced with *murM*$^{R6}$.** Cell wall of exponentially growing cultures ($OD_{550nm}$=0.4-0.5) was isolated, treated with LytA and the stem peptides were separated using C18 reverse phase HPLC. The stem petides of the numbered peaks are illustrated in **Figure S3**. The replacement of *murM* resulted in a shift from consisting of mostly branched stem peptides to a more Wt-like cell wall consistent with previous research (Filipe et al., 2000a).

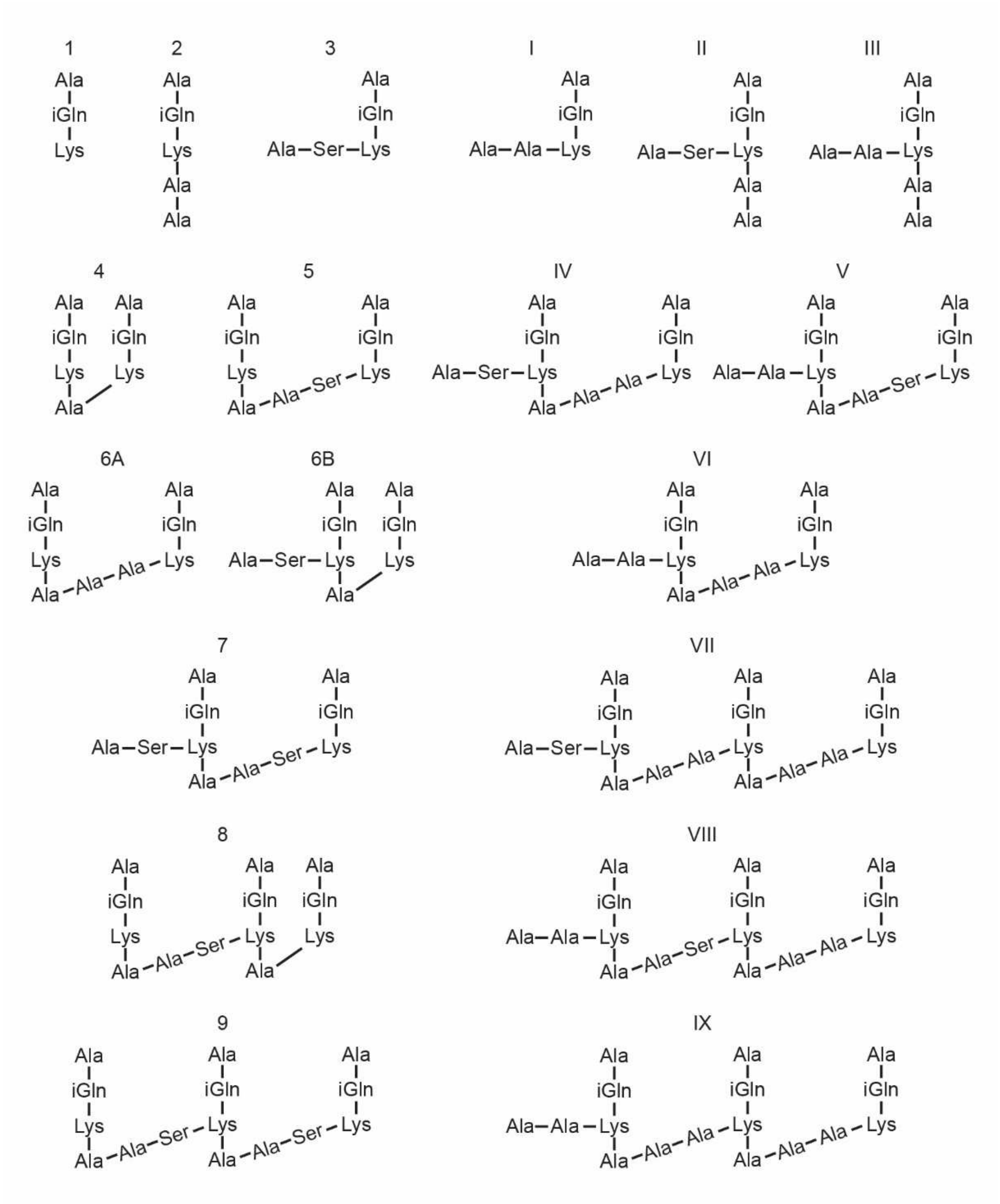


**Figure S3: Structures of pneumococcal cell wall stem peptides.** The stem peptides structures have been published previously (Garcia-Bustos et al., 1987; Garcia-Bustos et al., 1988; Severin & Tomasz, 1996).

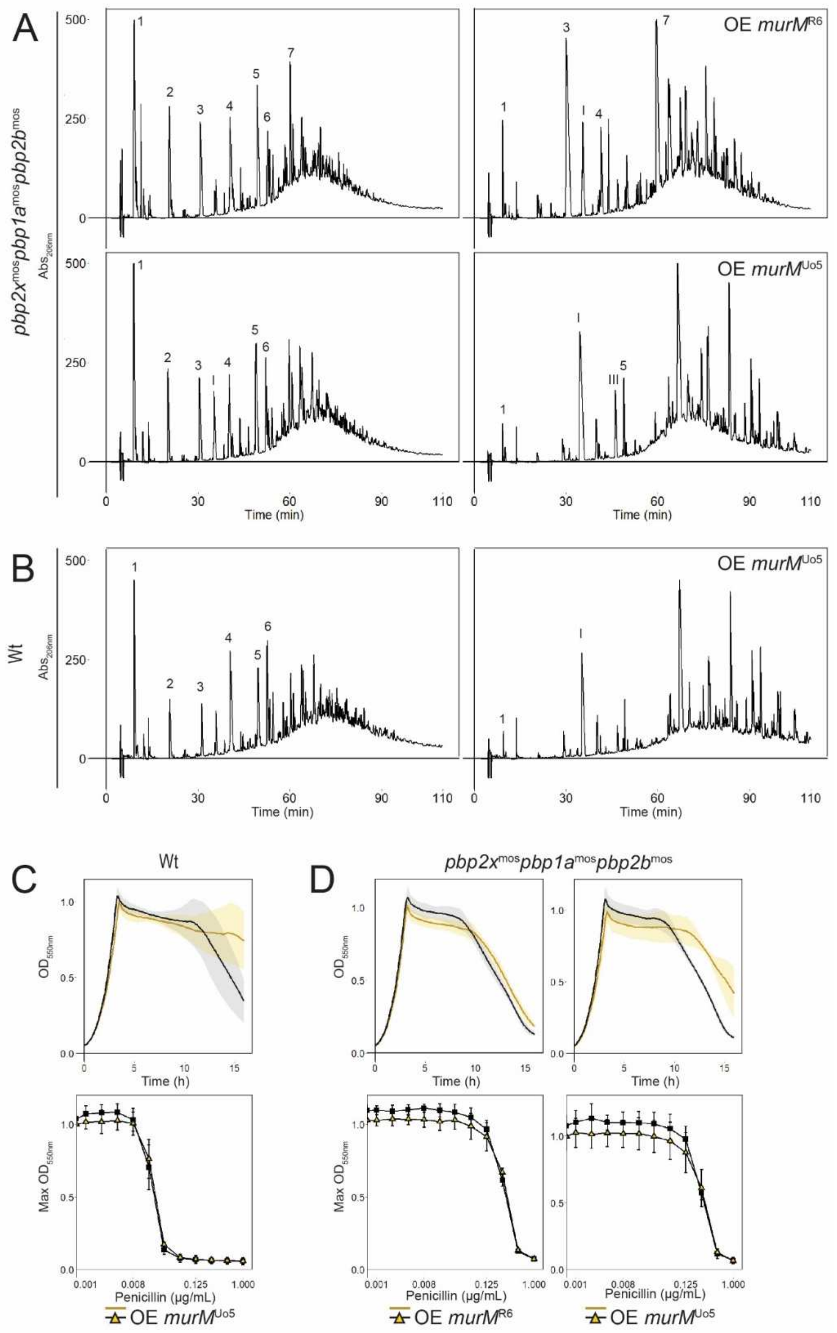


**Figure S4: Phenotypical characteristics of *murM* overexpression in Wt and low-affnity PBP background.** *murM* from either R6 or Uo5 (as indicated in the figure) was placed behind an inducible promoter ($P_{comX}$ using the ComRS system (Berg et al., 2011)) in Wt (**B**, **C**) or the *pbp2x*$^{mos}$*pbp1a*$^{mos}$*pbp2b*$^{mos}$ mutant (**A**, **D**). (**A**, **B**) Cultures were induced with ComS (0.2 µM) at $OD_{550nm}$ = 0.05 and cell wall of exponentially growing cultures ($OD_{550nm}$ = 0.4-0.5) was isolated, treated with LytA and the stem peptides were separated using C18 reverse phase HPLC. Stem peptide sturctures of the indicated peaks are illutrated in **Figure S3**. Overexpression (OE) of *murM* resulted in a clear shift from a linear to a more branched cell wall structure. (**C**, **D**) Cultures were diluted to an $OD_{550nm}$ of 0.05, inducer (0.2 µM ComS) was added, and bacterial growth ($OD_{550nm}$) was measured continuously every 5 minutes for 16 hours. The MIC curves display maximum $OD_{550nm}$ at each PenG concentration (a two-fold dilution series starting at 1µg/mL). The black lines and points represent control measurements (without inducer) and the yellow lines and points represent *murM* overexpression. Standard deviation was calculated from three biological replicates. Overexpression of *murM* had no effect on $MIC_{50}$ or growth rate but showed delayed autolysis.

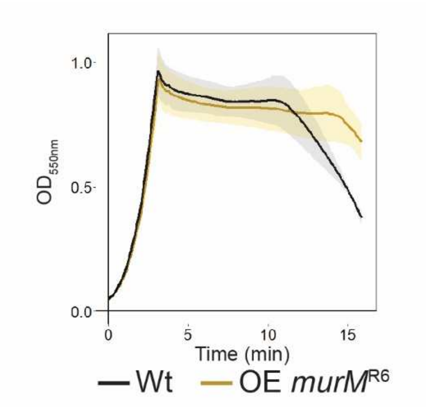


**Figure S5: Growth during *murM* overexpression in Wt.** *murM*$^{R6}$ was overexpressed (OE) in Wt using the ComRS system. Cultures were diluted to an $OD_{550nm}$ of 0.05, inducer (0.2 µM ComS) was added, and bacterial growth ($OD_{550nm}$) was measured continuously every 5 minutes for 16 hours. Error bars represent the standard deviation calculated from three biological replicates. Overexpression of *murM* had no effect on growth rate but showed delayed autolysis.

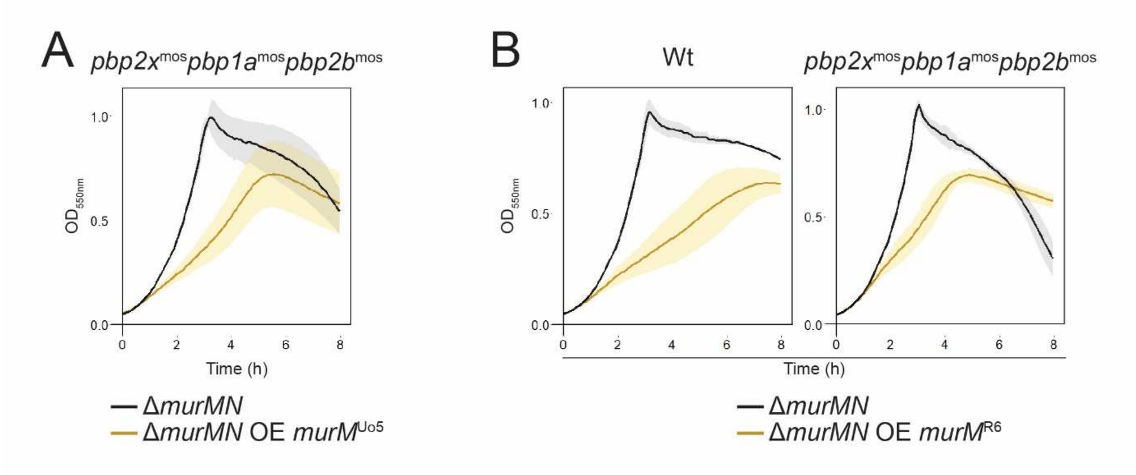


**Figure S6: Impact of *murM* overexpression in Δ*murMN* mutants.** *murM*$^{Uo5}$ (**A**) and *murM*$^{R6}$ (**B**) was overexpressed (OE) in Δ*murMN* mutants of Wt and *pbp2x*$^{mos}$*pbp1a*$^{mos}$*pbp2b*$^{mos}$ cells using the ComRS system. Cultures were diluted to an $OD_{550nm}$ of 0.05, inducer (0.2 µM ComS) was added and bacterial growth ($OD_{550nm}$) was measured continuously every 5 minutes for 16 hours (only the first 8h are displayed). Standard deviation was calculated from three biological replicates. A toxic effect of *murM* overexpression was observed when *murN* was deleted.

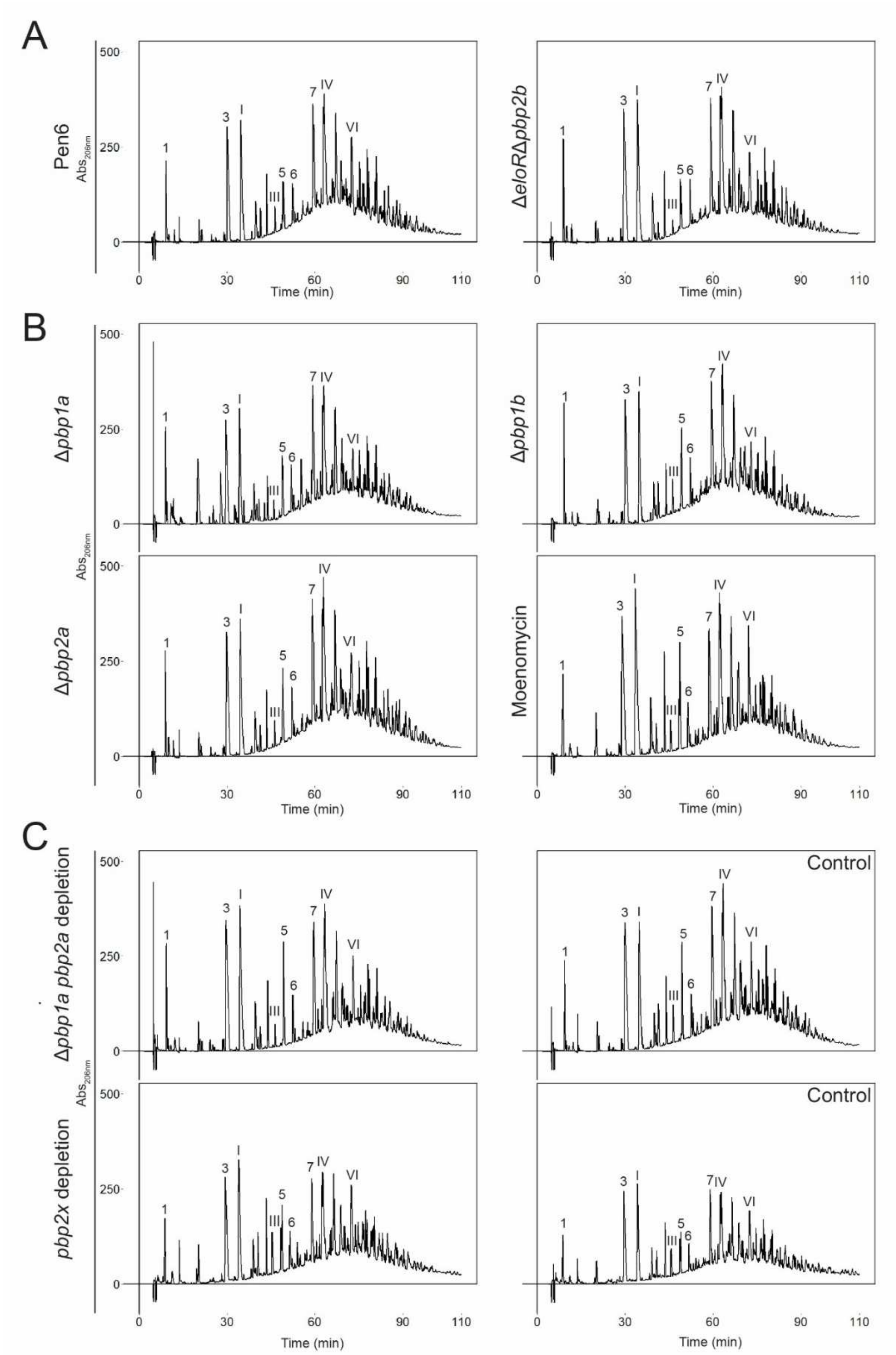


**Figure S7: Cell wall stem peptide composition of Pen6 *pbp* mutants.** Cell wall was isolated from cells at $OD_{550nm}$ = 0.4-0.5, treated with LytA and the stem peptides were separated using C18 reverse phase HPLC. Stem peptide structures of the indicated peaks are illutrated in **Figure S3**. The area under the peaks were quantified and percentage of each component are listed in **Table S1**. (**A**) The parental strain Pen6 and a double Δ*eloRΔpbp2b* mutant. No major differences were observed in the mutant. (**B**) Stem peptide profiles of single Class A *pbp*s mutants and moenomycin (5 µg/mL) treated cells. The moenomycin specifically targets the Class A PBPs and cell wall wals isolated from growth inhibited cells. Knockout or inhibition of class A PBPs had little influence on the cell wall composition. (**C**) Stem peptide profiles of cells depleted for *pbp2a* in a Δ*pbp1a* background and cells depleted of *pbp2x*. The *pbp*-depleted cells were harvested at the point where cells grew poorly from lack of the PBP, but were still viable. No evident changes to the stem peptide composition were observed.

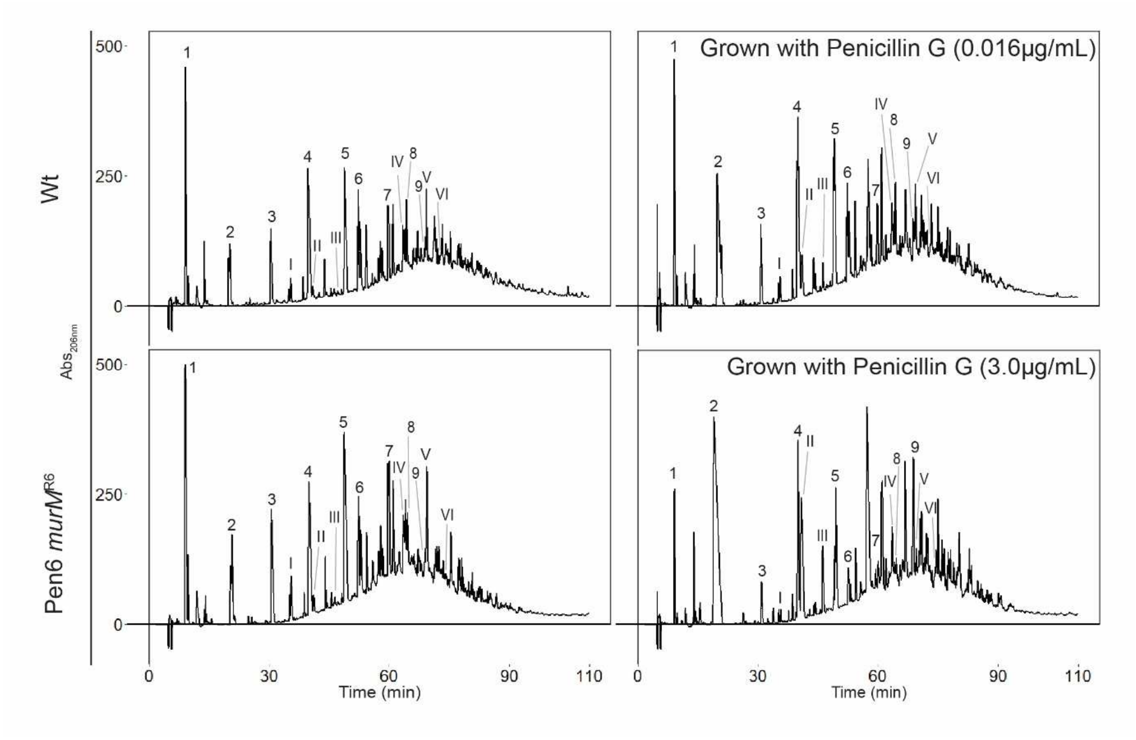


**Figure S8: Changes to the cell wall stem peptide composition during penicillin exposure.** Stem peptide profile of Wt and a Pen6 *murM*[R6] mutant when grown with subinhibitory concentrations of PenG (indicated) compared to non-treated controls. PenG was added to the growing culture at $OD_{550nm}$ = 0.05 and cell wall was isolated when the cultures reached $OD_{550nm}$ = 0.4-0.5. Stem peptide structures of the indicated peaks are illutrated in **Figure S3**. The area of the peaks of the stem peptides were quantified and percentage of each component are listed in **Table 2**. Upon PenG exposure, the peptidoglycan composition of both strains displayed a percentage incresae in monomeres as well as linear peptides relative to branched peptides.

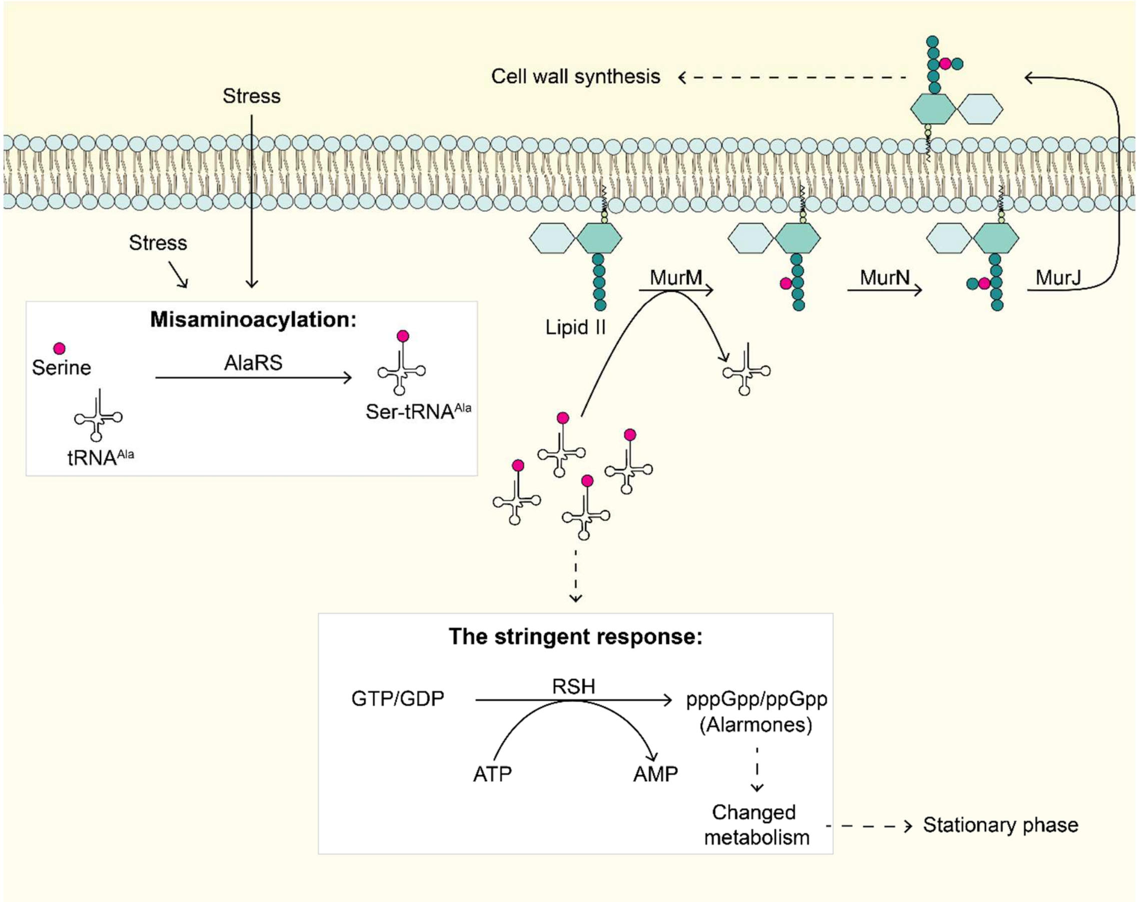


**Figure S9: The interpaly between the stringent response and MurM in pneumococci.** Alanyl-tRNA$^{Ala}$ synthetase (AlaRS) recognizes alanine and synthesizes Ala- tRNA$^{Ala}$ but it can also mistakenly recognize serine, leading to misaminoacylated Ser- tRNA$^{Ala}$. AlaRS has editing activity and can edit the misaminoacylated tRNAs, but the enzyme is error prone, leading to mismatched tRNAs. Stress (acidic stress was previously tested) leads to a higher number of misaminoacylated Ser-tRNA$^{Ala}$, which could lead to translation errors. A paper from Aggarwal et al. (2021) found that MurM is likely to work as a buffer for missaminoacylated Ser- tRNA$^{Ala}$ by favouring incorporation of these serine amino acids into the cell wall. Deletion of *murMN* under acidic stress led to accumulation of Ser- tRNA$^{Ala}$, activating the stringent response pathway. Activation of the stringent response triggers production of a large amount of alarmones (pppGpp or ppGpp) in the cells. In Gram positives, the alarmones are produced by the RSH protein (RelA/SpoT homolog) that has both synthesase and hydrolase activity. Increased alarmone levels leads to changes in the cell's metabolism and coordinate the entry into stationary phase.

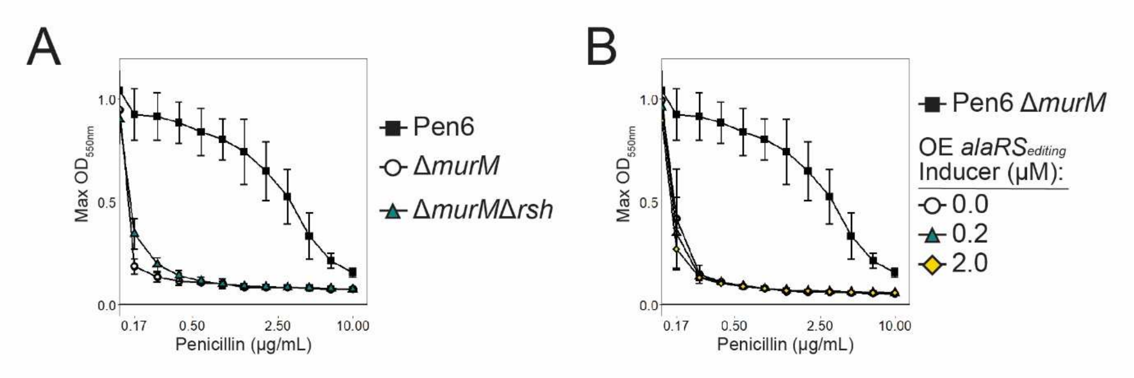


**Figure S10: Inhibition of the stringent response in Δ*murM* mutants.**. Panel **A** shows the PenG MIC curves of Pen6 and its Δ*murM* and a double Δ*murM*Δ*rsh* mutant. Deletion of *rsh* in a Δ*murM* mutant did not revert the initial phenotype of the strain. (**B**) The editing domain of AlaRS ($alaRS_{editing}$) was ectopically expressed using the ComRS system in a Pen6 Δ*murM* mutant. Overexpression (OE) of $AlaRS_{editing}$ using 2 µM ComS inducer gave no change in $MIC_{50}$ in the Δ*murM* mutant. Error bars represent standard deviation calculated from three biological replicates.

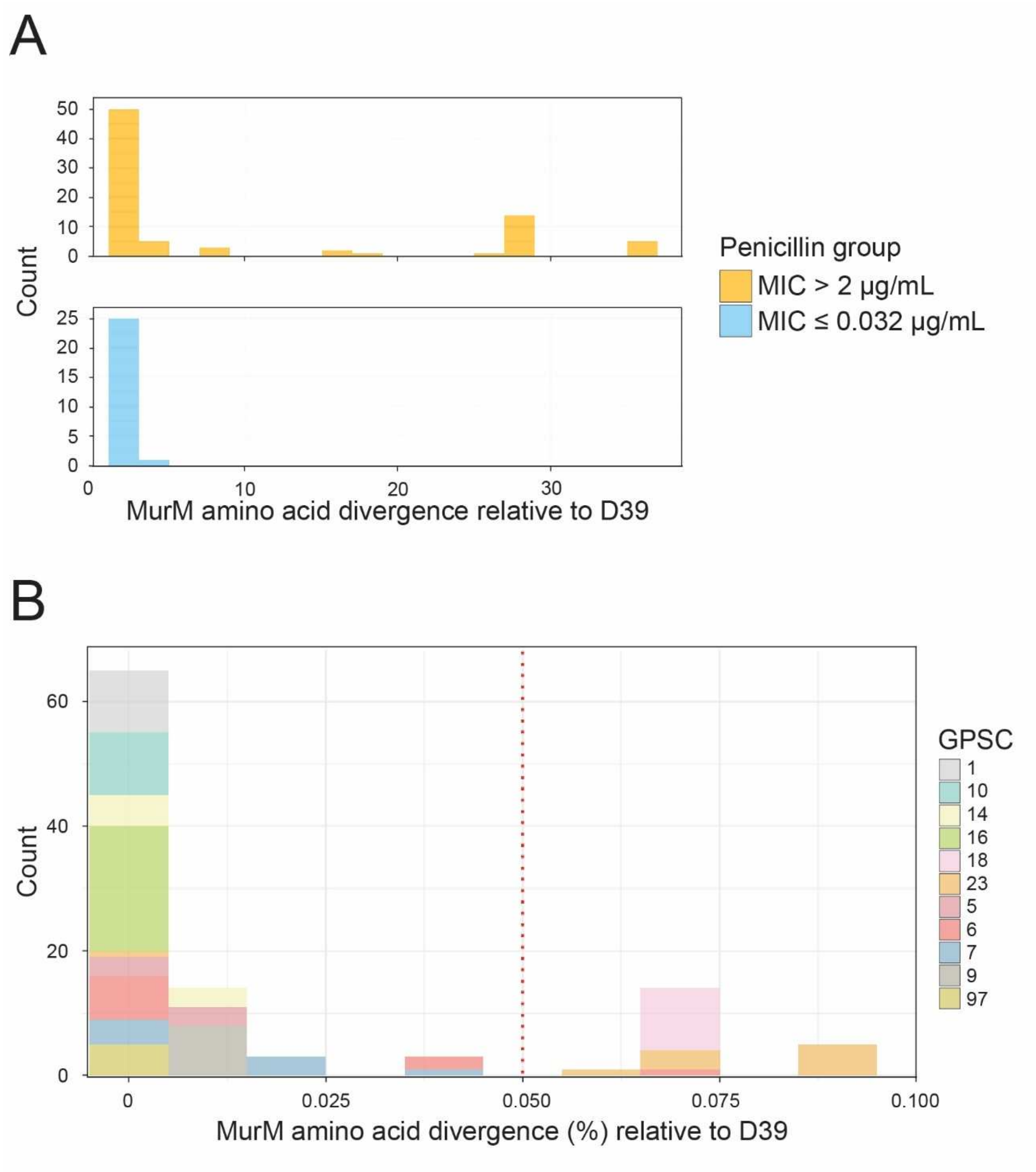


**Figure S11: MurM divergence and PenG MICs.** (**A**) Histogram of MurM divergence (amino acid differences) relative to the D39 reference, stratified by MIC groups (resistant isolates on top, susceptible bottom). (**B**) Histogram of MurM divergence (%) relative to the D39 reference, colored by GPSC type. The Wt used in this study (R6) is a non-encapsulated derivate of the strain D39 and have identical MurM sequences.

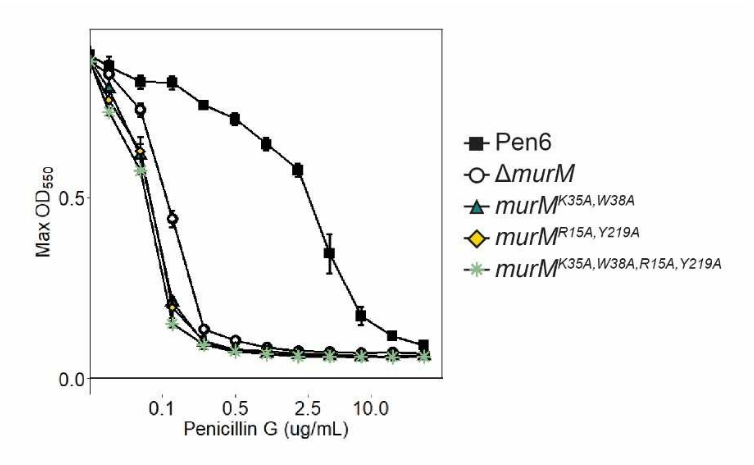


**Figure S12: Resistance profiles of catalytically inactive MurM mutants.** Curves display maximum $OD_{550nm}$ obtained at each PenG concentration. Standard deviation was calculated from three biological replicates. Presence of a non-functional MurM (i.e., mutants with essential amino acid recidues for lipid II binding replaced) did not rescue the Δ*murM* phenotype in Pen6.

**Supplementary Tables**

**Table S1:** Stem peptide composition of Pen6 and PBP inhibited or knockout mutants.

| | | | | | | | | | | | $\mathbf{P}_{comX}$-***pbp2a*. Δ*pbp2a*Δ*pbp1a*** | | $\mathbf{P}_{comX}$-***pbp2x*. Δ*pbp2x*** | |
|---|---|---|---|---|---|---|---|---|---|---|---|---|---|---|
| **Peak** | **Molecular Weight[a]** | **Peptide characteristics** | | **Pen6** | **Δ*pbp1b*** | **Δ*pbp2a*** | **Δ*pbp1a*** | **Δ*eloR*** | **Δ*eloR*Δ*pbp2b*** | **Moenomycin** | **Depletion** | **Ctr** | **Depletion** | **Ctr** |
| 1 | 345.2 | Linear | Monomer | 4.2 | 5.9 | 4.9 | 4.8 | 4.2 | 4.8 | 4.3 | 4.9 | 4.0 | 4.6 | 4.1 |
| 2 | 487.3 | Linear | Monomer | 1.6 | 1.3 | 1.4 | 7.6 | 1.2 | 1.2 | 2.3 | 1.6 | 1.9 | 1.0 | 2.4 |
| 3 | 503.4 | Branched | Monomer | 18.9 | 18.4 | 18.1 | 15.0 | 20.0 | 21.6 | 18.7 | 20.3 | 18.7 | 17.0 | 18.0 |
| I | 487.4 | Branched | Monomer | 16.7 | 17.0 | 17.3 | 13.8 | 19.2 | 20.3 | 21.1 | 21.6 | 15.6 | 16.6 | 16.0 |
| 4 | 743.4 | Linear | Dimer | 0.3 | 0.7 | 0.4 | 1.5 | 0.4 | 0.4 | 0.5 | 0.5 | 0.6 | 0.9 | 0.7 |
| II | 645.6 | Branched | Monomer | 1.6 | 2.1 | 1.3 | 1.1 | 1.5 | 0.8 | 1.3 | 1.1 | 2.4 | 3.2 | 2.4 |
| III | 629.4 | Branched | Monomer | 1.7 | 2.2 | 1.3 | 1.0 | 1.4 | 0.7 | 1.4 | 1.1 | 2.3 | 3.6 | 2.9 |
| 5 | 901.5 | Branched | Dimer | 6.1 | 6.5 | 6.1 | 6.5 | 5.1 | 4.8 | 7.0 | 6.4 | 5.6 | 7.0 | 6.2 |
| 6a | 885.5 | Branched | Dimer | 2.7 | 3.6 | 3.3 | 3.6 | 2.8 | 3.0 | 2.2 | 3.0 | 2.6 | 2.5 | 2.1 |
| 6b | 901.5 | Branched | Dimer | 0.5 | 0.8 | 0.5 | 0.7 | 0.5 | 0.4 | 0.4 | 0.5 | 1.0 | 0.5 | 0.3 |
| 7 | 1059.6 | Branched | Dimer | 7.5 | 9.1 | 8.8 | 11.9 | 7.4 | 7.6 | 8.5 | 8.8 | 10.2 | 9.3 | 11.1 |
| IV | 1044.3 | Branched | Dimer | 18.1 | 19.0 | 18.6 | 19.2 | 19.1 | 18.3 | 15.8 | 15.1 | 18.4 | 15.2 | 15.8 |
| 8 | 1300.5 | Branched | Trimer | 0 | 0 | 0 | 0 | 0 | 0 | 0 | 0 | 0 | 0 | 0 |
| 9 | 1458.6 | Branched | Trimer | 1.8 | 1.0 | 1.4 | 1.3 | 1.3 | 1.4 | 1.8 | 1.4 | 2.2 | 2.6 | 2.3 |
| V | 1044.3 | Branched | Dimer | 2.6 | 3.0 | 2.0 | 2.5 | 2.1 | 2.4 | 3.6 | 2.1 | 2.8 | 3.5 | 2.7 |
| VI | 1028.1 | Branched | Dimer | 8.7 | 4.9 | 7.0 | 4.6 | 7.0 | 6.6 | 6.6 | 5.8 | 6.8 | 7.3 | 7.5 |
| VII | 1584.9 | Branched | Trimer | 4.1 | 2.6 | 4.3 | 3.0 | 4.2 | 3.3 | 2.2 | 3.3 | 2.6 | 2.2 | 2.4 |
| VIII | 1584.9 | Branched | Trimer | 2.0 | 1.3 | 2.0 | 1.4 | 1.7 | 1.7 | 1.5 | 1.9 | 1.8 | 2.2 | 2.2 |
| IX | 1568.9 | Branched | Trimer | 0.8 | 0.5 | 1.1 | 0.5 | 1.0 | 0.8 | 0.6 | 0.4 | 0.3 | 0.9 | 0.8 |
| **Total** | | | | 100 | 100 | 100 | 100 | 100 | 100 | 100 | 100 | 100 | 100 | 100 |
| **Monomers (%)** | | | | 44.9 | 47.0 | 44.4 | 43.2 | 47.6 | 49.4 | 49.2 | 50.7 | 45.0 | 46.0 | 45.9 |
| **Oligomers (%)** | | | | 55.1 | 53.0 | 55.6 | 56.8 | 52.4 | 50.6 | 50.8 | 49.3 | 55.0 | 54.0 | 54.1 |
| **Linear peptides (%)** | | | | 6.1 | 8.0 | 6.7 | 13.8 | 5.8 | 6.4 | 7.2 | 7.0 | 6.5 | 6.5 | 7.2 |
| **Branched peptides (%)** | | | | 93.9 | 92.0 | 93.3 | 86.2 | 94.2 | 93.6 | 92.8 | 93.0 | 93.5 | 93.5 | 92.8 |
| **B/L peptides** | | | | 15.3 | 11.5 | 14.0 | 6.2 | 16.2 | 14.7 | 13.0 | 13.2 | 14.3 | 14.5 | 12.8 |

[a]Stem peptides were identified based on molecular mass published in previous literature (Garcia-Bustos et al., 1987; Garcia-Bustos et al., 1988; Severin & Tomasz, 1996)

**Table S2:** Strains and mutants used in this work.

| Strains | Relevant characteristics | Reference |
|---|---|---|
| ***Streptococcus oralis*:** | | |
| Uo5 | A high level β-lactam resistant isolate of *S. oralis* isolated from a Hungarian nasal swab | Reichmann et al. (1997) |
| ***Streptococcus pneumoniae*:** | | |
| RH425 | R6 derivative, but Δ*comA*::*ermAM*, *rpsL1*; Ery$^r$ , Sm$^r$ | Johnsborg and Håvarstein (2009) |
| SPH131 | RH425, but Δ*IS1167*::*P1-comR*, Δ*cps*::P$_{comX}$-janus; Ery$^r$ , Kan$^r$ | Berg et al. (2011) |
| Pen6 | R6Hex transformant with chromosomal DNA from penicillin-resistant clinical isolate 8249 and selected for Pen$^r$ | Zighelboim and Tomasz (1980) |
| MH10 | RH425, but *pbp2*x$^{mos}$; Ery$^r$ , Sm$^r$ | This work |
| MH56 | RH425, but *pbp2x*$^{mos}$*pbp1a*$^{mos}$; Ery$^r$ , Sm$^r$ | This work |
| MH68 | RH425, but *pbp2x*$^{mos}$*pbp2b*$^{mos}$; Ery$^r$ , Sm$^r$ | This work |
| MH83 | RH425, but *pbp2x*$^{mos}$*pbp1a*$^{mos}$*pbp2b*$^{mos}$; Ery$^r$ , Sm$^r$ | This work |
| MH105 | RH425, but Δ*IS1167*::*P1-comR*, Δ*cps*::P$_{comX}$ -*murM*$^{Uo5}$; Ery$^r$ , Sm$^r$ | This work |
| MH130 | RH425, but *pbp2x*$^{mos}$*pbp1a*$^{mos}$*pbp2b*$^{mos}$, Δ*IS1167*::janus; Ery$^r$ , Kan$^r$ | This work |
| MH132 | RH425, but *pbp2x*$^{mos}$*pbp1a*$^{mos}$*pbp2b*$^{mos}$, Δ*IS1167*::*P1-comR*; Ery$^r$ , Sm$^r$ | This work |
| MH134 | RH425, but *pbp2x*$^{mos}$*pbp1a*$^{mos}$*pbp2b*$^{mos}$, Δ*IS1167*::*P1-comR*, Δ*cps*::P$_{comX}$ -janus; Ery$^r$ , Kan$^r$ | This work |
| MH136 | RH425, but Δ*IS1167*::*P1-comR*, Δ*cps*::P$_{comX}$ -*murM*$^{Uo5}$, Δ*murMN*::janus; Ery$^r$ , Kan$^r$ | This work |
| MH138 | RH425, but *pbp2x*$^{mos}$*pbp1a*$^{mos}$*pbp2b*$^{mos}$, Δ*IS1167*::*P1-comR*, Δ*cps*::P$_{comX}$ -*murM*$^{Uo5}$; Ery$^r$ , Sm$^r$ | This work |
| MH141 | RH425, but *pbp2x*$^{mos}$*pbp1a*$^{mos}$*pbp2b*$^{mos}$, Δ*IS1167*::*P1-comR*, Δ*cps*::P$_{comX}$ -*murM*$^{Uo5}$, Δ*murMN*::janus; Ery$^r$ , Kan$^r$ | This work |
| MH144 | RH425, but *pbp2x*$^{mos}$*pbp1a*$^{mos}$*pbp2b*$^{mos}$, Δ*IS1167*::*P1-comR*, Δ*cps*::P$_{comX}$ -*murM*$^{R6}$; Ery$^r$ , Sm$^r$ | This work |
| MH146 | RH425, but Δ*is1167*::*P1-comR*, Δ*cps*::P$_{comX}$ -*murM*$^{R6}$, Δ*murMN*::janus; Ery$^r$ , Kan$^r$ | This work |
| MH147 | RH425, but *pbp2x*$^{mos}$*pbp1a*$^{mos}$*pbp2b*$^{mos}$, Δ*IS1167*::*P1-comR*, Δ*cps*::P$_{comX}$ -*murM*$^{R6}$, Δ*murMN*::janus; Ery$^r$ , Kan$^r$ | This work |
| MH149 | RH425, but Δ*IS1167*::*P1-comR*, Δ*cps*::P$_{comX}$ -*murM*$^{Uo5}$, Δ*murN*::janus; Ery$^r$ , Kan$^r$ | This work |
| RSG 66 | RH425, but Δ*murM*::janus; Ery$^r$ , Kan$^r$ | This work |
| RSG 69 | RH425, but *pbp2x*$^{mos}$*pbp1a*$^{mos}$*pbp2b*$^{mos}$, Δ*murM*::janus; Ery$^r$ , Kan$^r$ | This work |
| RSG73 | RH425, but *murM*$^{Uo5}$; Ery$^r$ , Sm$^r$ | This work |
| RSG75 | RH425, but *pbp2x*$^{mos}$*pbp1a*$^{mos}$*pbp2b*$^{mos}$, *murM*$^{Uo5}$; Ery$^r$ , Sm$^r$ | This work |
| RSG173 | Pen6, but Sm$^r$ | This work |
| RSG183 | Pen6, but Δ*murM*::janus; Kan$^r$ | This work |
| RSG185 | Pen6, but Δ*eloR*::janus; Kan$^r$ | This work |
| RSG186 | Pen6, but *murM*$^{R6}$; Sm$^r$ | This work |
| RSG189 | Pen6, but Δ*eloR*::DEL; Sm$^r$ | This work |
| RSG192 | Pen6, but Δ*IS1167*::janus;Kan$^r$ | This work |
| RSG194 | Pen6, but Δ*IS1167*::*P1-comR*; Sm$^r$ | This work |
| RSG199 | Pen6, but Δ*IS1167*::*P1-comR*, Δ*cps*::P$_{comX}$ -janus; Kan$^r$ | This work |
| RSG200 | RH425, but Δ*IS1167*::*P1-comR*, Δ*cps*::P$_{comX}$ -*murM*$^{R6}$; Sm$^r$ | This work |
| RSG203 | Pen6, but Δ*IS1167*::*P1-comR*, Δ*cps*::P$_{comX}$ -*pbp2a*; Sm$^r$ | This work |

| | | |
|---|---|---|
| RSG206 | Pen6, but Δ*IS1167*::*P1-comR*, Δ*cps*::$P_{comX}$-*pbp2a*, Δ*pbp2a*::janus; $Kan^r$ | This work |
| RSG207 | Pen6, but Δ*is1167*::*P1-comR*, Δ*cps*::$P_{comX}$-*pbp2a*, Δ*pbp2a*::DEL; $Sm^r$ | This work |
| RSG208 | Pen6, but Δ*is1167*::*P1-comR*, Δ*cps*::$P_{comX}$-*pbp2a*, Δ*pbp2a*::DEL, Δ*pbp1a*::janus; $Kan^r$ | This work |
| RSG214 | Pen6, but Δ*murM*::DEL; $Sm^r$ | This work |
| RSG219 | Pen6, but Δ*murM*::DEL, Δ*rsh*::janus; $Kan^r$ | This work |
| RSG234 | RH425, but Δ*is1167*::*P1-comR*, Δ*cps*::$P_{comX}$-$murM^{R6}$, Δ*murM*::janus; $Kan^r$ | This work |
| RSG235 | Pen6, but Δ*is1167*::*P1-comR*, Δ*cps*::$P_{comX}$-$alaRS_{editing}$; $Sm^r$ | This work |
| RSG243 | Pen6, but Δ*is1167*::*P1-comR*, Δ*cps*::$P_{comX}$-$alaRS_{editing}$, Δ*murM*::janus; $Kan^r$ | This work |
| JM5 | Pen6, but Δ*pbp1b*::janus; $Kan^r$ | This work |
| JM6 | Pen6, but Δ*pbp2a*::janus; $Kan^r$ | This work |
| JM9 | Pen6, but Δ*pbp1a*::janus; $Kan^r$ | This work |
| JM12 | Pen6, but Δ*eloR*::DEL, Δ*pbp2b*::janus; $Kan^r$ | This work |
| AW520 | Pen6, but Δ*is1167*::*P1-comR*, Δ*cps*::$P_{comX}$-$pbp2x^{Pen6}$; $Sm^r$ | This work |
| AW524 | Pen6, but Δ*is1167*::*P1-comR*, Δ*cps*::$P_{comX}$-$pbp2x^{Pen6}$, Δ*pbp2x*::janus; $Kan^r$ | This work |
| AW594 | Pen6, but Δ*is1167*::*P1-comR*, Δ*cps*::$P_{comX}$-$pbp2x^{Pen6}$, Δ*pbp2x*::janus, Δ*lytA*::*aad9*; $Sm^r$, $Spc^r$ | This work |
| AW627 | Pen6, but $murM^{K35A, W38A}$; $Sm^r$ | This work |
| AW628 | Pen6, but $murM^{R15A, Y219A}$; $Sm^r$ | This work |
| AW629 | Pen6, but $murM^{K35A, W38A, R15A, Y219A}$; $Sm^r$ | This work |

**Table S3:** Primers used in this work

| Primer | Description | Sequence(5'-->3') | Reference |
|---|---|---|---|
| **Primers to amplify the *pbp1a*$^{\mathrm{Uo5}}$ amplicon and sequencing of the mutants** | | | |
| MVH26 | ~1000bp upstream *pbp1a*$^{\mathrm{Uo5}}$ | CCCTTGTGCTCATATTGTGG | This work |
| MVH27 | ~1000bp downstream *pbp1a*$^{\mathrm{Uo5}}$ | TCTGAGCCAACTAATGCCAAC | This work |
| MVH40 | ~500 bp in *pbp1a*$^{\mathrm{Uo5}}$ | AGAGATCTTGACCTACTAC | This work |
| MVH41 | ~1000 bp in *pbp1a*$^{\mathrm{Uo5}}$ | TCATTGCTCAGTTAGGTTCTCG | This work |
| MVH42 | ~1500 bp in *pbp1a*$^{\mathrm{Uo5}}$ | GTATTTAGTGATGGTAGC | This work |
| **Primers to amplify the *pbp2b*$^{\mathrm{Uo5}}$ amplicon and sequencing of the mutants** | | | |
| MVH24 | ~1000bp upstream *pbp2b*$^{\mathrm{Uo5}}$ | AGGCATAAATCAAATCTATTAAAATG | This work |
| MVH25 | ~1000bp downstream *pbp2b*$^{\mathrm{Uo5}}$ | TGATTTTGCTTCTTGCTCGTG | This work |
| MVH37 | ~500 bp in *pbp2b*$^{\mathrm{Uo5}}$ | CTATCTCTTTAGCCAGCTCAATG | This work |
| MVH38 | ~1000 bp in *pbp2b*$^{\mathrm{Uo5}}$ | CTGAAGGTGTCTATGCAGTAG | This work |
| MVH39 | ~1500 bp in *pbp2b*$^{\mathrm{Uo5}}$ | GCCAGTTTGATAACTACACACC | This work |
| **Primers to amplify the *pbp2x*$^{\mathrm{Uo5}}$ amplicon and sequencing of the mutants** | | | |
| MVH22 | ~1000bp upstream *pbp2x*$^{\mathrm{Uo5}}$ | TGGTGTCCAGGAAATTGATGG | This work |
| MVH23 | ~1000bp downstream *pbp2x*$^{\mathrm{Uo5}}$ | TGTAATCAAAAGTTAGTTTTACAG | This work |
| MVH34 | ~500 bp in *pbp2x*$^{\mathrm{Uo5}}$ | GATGTCCATTAAACAAGAC | This work |
| MVH35 | ~1000 bp in *pbp2x*$^{\mathrm{Uo5}}$ | GGATCAGGCATGAAGGTTATG | This work |
| MVH36 | ~1500 bp in *pbp2x*$^{\mathrm{Uo5}}$ | CACATGATCTTAGTTGGGACG | This work |
| **Primers to amplify *murM* and *murN* sequences** | | | |
| VE47 | ~1000bp upstream *murM*$^{\mathrm{R6}}$ | ACCAGTAGTCATGGAAGCAAA | (Berg et al., 2013) |
| KHB199 | ~1000bp downstream *murN*$^{\mathrm{R6}}$ | CACAATTTCAGACACCAGAGC | (Straume et al., 2020) |
| MVH43 | ~100 bp upstream *murM*$^{\mathrm{R6}}$ (also ~1300bp upstream *murN*$^{\mathrm{R6}}$) | CTTAGTTTGAACTTCAGCATAG | This work |
| MVH44 | ~100 bp downstream *murN*$^{\mathrm{R6}}$ (also ~1300bp downstream *murM*$^{\mathrm{R6}}$) | GCCAGCGCATGTCTCTCC | This work |
| MVH45 | ~100 bp downstream *murM*$^{\mathrm{R6}}$ | CTAGCAAATCCCCCATCTGG | This work |
| **Primers to amplify the Janus cassette** | | | |
| Kan484F | Start Janus cassette | GTTTGATTTTTAATGGATAATGTG | (Johnsborg et al., 2008) |
| RpsL41R | End Janus cassette | CTTTCCTTATGCTTTTGGAC | (Johnsborg et al., 2008) |
| **Primers to create the Δ*murMN*::janus amplicon** | | | |
| VE47 | ~1000bp upstream *murM*$^{\mathrm{R6}}$ | ACCAGTAGTCATGGAAGCAAA | (Berg et al., 2013) |
| KHB199 | ~1000bp downstream *murN*$^{\mathrm{R6}}$ | CACAATTTCAGACACCAGAGC | (Straume et al., 2020) |
| | Template strain: MH110 (Δ*comA*, Δ*murMN*::janus, Ery$^{r}$ Kan$^{r}$) | | (Straume et al., 2020) |
| **Primers to create the *ΔmurN*::janus amplicon** | | | |
| MVH43 | ~100 bp upstream *murM*$^{\mathrm{R6}}$ (also ~1300bp upstream *murN*$^{\mathrm{R6}}$) | CTTAGTTTGAACTTCAGCATAG | This work |
| KHB199 | ~1000bp downstream *murN*$^{\mathrm{R6}}$ | CACAATTTCAGACACCAGAGC | (Straume et al., 2020) |
| KHB198 | End Janus cassette, **overlap just down *murN*$^{\mathrm{R6}}$** | CTAAACGTCCAAAAGCATAAGGAAAG**GATGAAAAAGTCAGTATTTAGATT** | (Straume et al., 2020) |
| MVH49 | start Janus cassette, **overlap start *murN*$^{\mathrm{R6}}$** | CACATTATCCATTAAAAATCAAAC*CTTC***TTTCGTGAGTGTTGTTAG** | This work |

| | | | |
|---|---|---|---|
| **Template to amplify the Δ*murM*::janus amplicon** | | | |
| VE47 | ~1000bp upstream *murM*$^{R6}$ | ACCAGTAGTCATGGAAGCAAA | (Berg et al., 2013) |
| MVH44 | ~100 bp downstream *murN*$^{R6}$ (also ~1300bp downstream *murM*$^{R6}$) | GCCAGCGCATGTCTCTCC | This work |
| | Template strain: SPH181 (Δ*comA P1*::*PcomR*::*comR PcomX*::*pbp2b* Δ*pbp2b*$^{Wt}$ Δ*lytA*::Spc$^{r}$ Δ*murM*::Janus Ery$^{r}$ Spc$^{r}$ Kan$^{r}$) | | (Berg et al., 2013) |
| **Primers to replace murMNR6 with murMUo5** | | | |
| GS428 | just downstream *murN*$^{R6}$ | GATGAAAAAGTCAGTATTTAGATT | This work |
| GS429 | just upstream *murM*$^{R6}$ | TTCCTACTCTCTTTCCTCCA | This work |
| GS430 | just upstream *murM*$^{R6}$, **overlap start *murM*$^{Uo5}$** | TGGAGGAAAGAGAGTAGGAA**ATGTTTACGTATAAAATGAATGTTG** | This work |
| GS431 | just downstream *murN*$^{R6}$, **overlap end *murM*$^{Uo5}$** | AATCTAAATACTGACTTTTTCATC**CTAATTCCTACTTCGAAGTTTC** | This work |
| **Primers to amplify the Δ*is1167*::janus and Δ*is1167*::*P1-comR* amplicons** | | | |
| AmiF | ~1000bp upstream *P1-comR* | CGGTGAAGGAAGTAAGAAGTTT | (Johnsborg & Håvarstein, 2009) |
| TreR | ~1000bp downstream *P1-comR* | GTGACGGCAGTCACATTCTC | (Johnsborg & Håvarstein, 2009) |
| | Template strain: RH426 (RH425, but Δ*IS1167*::Janus; Ery$^{r}$ Kan$^{r}$) | | (Johnsborg & Håvarstein, 2009) |
| | Template strain: SPH131 (RH425, but Δ*is1167*::*P1-comR*, Δ*cps*::*PcomX*::janus) | | (Berg et al., 2011) |
| **Primers to amplify the *PcomX*::janus amplicon and create replacements** | | | |
| KHB31 | ~800bp upstream *PcomX* | ATAACAAATCCAGTAGCTTTGG | (Berg et al., 2011) |
| KHB34 | ~800bp downstream *PcomX*::janus | CATCGGAACCTATACTCTTTTAG | (Berg et al., 2011) |
| KHB33 | just down *PcomX*::janus | TTTCTAATATGTAACTCTTCCCAAT | (Berg et al., 2011) |
| KHB36 | end of *PcomX* | TGAACCTCCAATAATAAATATAAAT | (Berg et al., 2011) |
| | Template strain: SPH131 (RH425, but Δ*is1167*::*P1-comR*, Δ*cps*::*PcomX*::janus) | | |
| **Primers to create the *PcomX*::*murM*$^{Uo5}$ amplicon** | | | |
| KHB31 | ~800bp upstream *PcomX* | ATAACAAATCCAGTAGCTTTGG | (Berg et al., 2011) |
| KHB34 | ~800bp downstream *PcomX*::janus | CATCGGAACCTATACTCTTTTAG | (Berg et al., 2011) |
| MVH46 | end of *PcomX*, **overlap start *murM*$^{Uo5}$** | ATTTATATTTATTATTGGAGGTTCA**ATGTTTACGTATAAAATGAATGTTG** | This work |
| MVH47 | just downstream *PcomX*::janus, **overlap end *murM*$^{Uo5}$** | ATTGGGAAGAGTTACATATTAGAAA**CTAATTCCTACTTCGAAGTTTC** | This work |
| **Primers to create the *PcomX*::*murM*$^{R6}$ amplicon** | | | |
| KHB31 | ~800bp upstream *PcomX* | ATAACAAATCCAGTAGCTTTGG | (Berg et al., 2011) |
| KHB34 | ~800bp downstream *PcomX*::janus | CATCGGAACCTATACTCTTTTAG | (Berg et al., 2011) |
| KHB374 | end of *PcomX*, **overlap start *murM*$^{R6}$** | ATTTATATTTATTATTGGAGGTTC**AATGTACCGTTATCAAATTGGCAT** | This work |
| KHB375 | just downstream *PcomX*::janus, **overlap end *murM*$^{R6}$** | ATTGGGAAGAGTTACATATTAGAAA**TTACTTTCTATGTTTTTTCTTAATG** | This work |
| **Primers to create the *murM*::DEL amplicon** | | | |

| | | | |
|---|---|---|---|
| RSG46 | Just downstream *murM*$^{R6}$, **overlapp just up *murM*$^{R6}$** | GAGTGTTGTTAGTGCCATATAC**TTCCTACTCTCTTTCCTCCAG** | This work |
| RSG60 | Just downstream *murM*$^{R6}$ | GTATATGGCACTAACAACACTC | This work |
| **Primers to amplify the Δ*eloR*::janus and create the Δ*eloR*::DEL amplicon** | | | |
| DS374 | ~900bp upstream *eloR* | CGAAACCTTGGGATACGCAG | (Stamsås et al., 2017) |
| DS377 | ~1000bp downstream *eloR* | CAGCACCCACGTTAAGCAAC | (Stamsås et al., 2017) |
| DS390 | Just downtream *eloR***, overlap just up *eloR*** | GAAATAAATAAGGAGGAATCTGGTA**GTAAAATCAGGTTTATCCTGATTTTTTGCTAG** | This work |
| DS378 | Just up *eloR* | TACCAGATTCCTCCTTATTTATTTC | This work |
| | Template strain: SPH472 (Δ*comA*, Δ*eloR*::janus, m(sf)GFP-*mltG*; Ery$^r$, Kan$^r$) | | (Stamsås et al., 2017) |
| **Primers to amplify the Δ*pbp2b*::janus amplicon** | | | |
| KHB129 | ~900bp upstream *pbp2b*$^{R6}$ | CGATAAAGAAGAGCATAGGAAG | (Berg et al., 2013) |
| KHB132 | ~1000bp downstream *pbp2b*$^{R6}$ | TCCCAATCAATGGTTTCATTGG | (Berg et al., 2013) |
| | Template strain: SPH156 (Δ*pbp2b*::janus, *PcomX*::*pbp2b*, Kan$^r$) | | (Berg et al., 2013) |
| **Primers to amplify the Δ*pbp1a*::janus amplicon** | | | |
| MTS5F | ~1000bp upstream *pbp1a*$^{R6}$ | CCTTGTGTTCATAGCGAGG | (Straume et al., 2017) |
| MTS8R | ~1000bp downstream *pbp1a*$^{R6}$ | AAAACGGCTTTGGTAGCAGATG | (Straume et al., 2017) |
| | Template strain: SPH344 (Δ*comA*, *ssbB*::*luc*, Δ*pbp1a*::Janus, Ery$^r$, Cm$^r$, Kan$^r$) | | (Straume et al., 2017) |
| **Primers to amplify the Δ*pbp1b*::janus amplicon** | | | |
| MTS9F | ~1200bp upstream *pbp1b*$^{R6}$ | GCCTGTACTTGGTAGTTTGG | (Straume et al., 2017) |
| MTS12R | ~1000bp downstream *pbp1b*$^{R6}$ | GACTATTCCAGTATAGCAC | (Straume et al., 2017) |
| | Template strain: SPH345 (Δ*comA*, *ssbB*::*luc*, Δ*pbp1b*::Janus, Ery$^r$, Cm$^r$, Kan$^r$) | | (Straume et al., 2017) |
| **Primers to amplify the Δ*pbp2a*::janus amplicon** | | | |
| MTS1F | ~1000bp upstream *pbp2a*$^{R6}$ | GCACAACTTGTTCGTACTCTTG | (Straume et al., 2017) |
| MTS4R | ~1000bp downstream *pbp2a*$^{R6}$ | AGGTTTACTTCTGCAACTGTG | (Straume et al., 2017) |
| | Template strain: SPH346 (Δ*comA*, *ssbB*::*luc*, Δ*pbp2a*::Janus, Ery$^r$, Cm$^r$, Kan$^r$) | | (Straume et al., 2017) |
| **template to amplify the *PcomX*::*pbp2a* amplicon** | | | |
| KHB31 | ~800bp upstream *PcomX* | ATAACAAATCCAGTAGCTTTGG | (Berg et al., 2011) |
| KHB34 | ~800bp downstream *PcomX*::janus | CATCGGAACCTATACTCTTTTAG | (Berg et al., 2011) |
| MTS17F | end of *PcomX*, **overlap start *pbp2a*** | ATTTATATTTATTATTGGAGGTTCA**ATGAAATTAGATAAATTATTTGAGAA** | This work |
| MTS18 | just downstream *PcomX*::janus, **overlap end *pbp2a*** | GGGAAGAGTTACATATTAGAAA**TTAGCGAAATAGATTGACTATCG** | This work |
| **Primers to create the Δ*pbp2a*::DEL amplicon** | | | |
| MTS1F | ~1000bp upstream *pbp2a*$^{R6}$ | GCACAACTTGTTCGTACTCTTG | (Straume et al., 2017) |
| MTS4R | ~1000bp downstream *pbp2a*$^{R6}$ | AGGTTTACTTCTGCAACTGTG | (Straume et al., 2017) |
| MTS15R | Just downstream *pbp2a*, **overlapp just up *pbp2a*** | GCTAGGCTTTGACAAGCATC**GCGTTTATTTTATCATCTTCATC** | This work |
| MTS16F | Just upstream *pbp2a*, **overlapp just down *pbp2a*** | GATGAAGATGATAAAATAAACGC**GATGCTTGTCAAAGCCTAGC** | This work |
| **Primers to amplify the *rpsL* gene to generate a Sm$^r$ resistant strain** | | | |

| | | | |
|---|---|---|---|
| DS827 | ~1000bp upstream *rpsL* | CATCTAGGTAATAGCCGTAGTC | This work |
| DS828 | ~1000bp downstream *rpsL* | GGCATCGACGTGAGCCATG | This work |
| | Template strain: RH425 (R6 derivative, but Δ*comA*::*ermAM*, *rpsL1*; $Ery^r$ , $Sm^r$) | | (Johnsborg & Håvarstein, 2009) |
| **Primers to create the Δ*pbp2x*::janus amplicon** | | | |
| KHB104 | ~700bp upstream *pbp2x*$^{R6}$ | GAAGTGAAGCCGATTGAGAC | (Berg et al., 2013) |
| KHB107 | ~700bp downstream *pbp2x*$^{R6}$ | ACACAATTCCGATAATCAAGAG | (Berg et al., 2013) |
| AW348 | ~100bp upstream *pbp2x*$^{Pen6}$ | TTGGCACCCTATATCGAAAAAG | This work |
| AW349 | Start Janus cassette, **overlap just up *pbp2x*$^{Pen6}$** | CACATTATCCATTAAAAATCAAAC**TCCGCTATTCGAATATTTTCATTG** | This work |
| AW350 | End Janus cassette, **overlap just down *pbp2x*$^{Pen6}$** | GTCCAAAAGCATAAGGAAAG**ATGTTTATTTCCATCAGTGCTGG** | This work |
| AW351 | ~100bp downstream *pbp2x*$^{Pen6}$ | TACTATATTTTGAGCAGCCTAAAG | This work |
| **Primers to create the *PcomX*::*pbp2x*$^{Pen6}$ amplicon** | | | |
| AW354 | end of *PcomX*, overlap start *pbp2x*$^{Pen6}$ | ATTTATATTTATTATTGGAGGTTCA**ATGAAGTGGACAAAAAGAATAACC** | This work |
| AW355 | Just downstream *PcomX*, overlap end *pbp2x*$^{Pen6}$ | ATTGGGAAGAGTTACATATTAGAAA**CAGCACTGATGGAAATAAACATATTA** | This work |
| AW356 | Sequencing primer ~700 bp into *pbp2x*$^{Pen6}$ | CGTCTGGGTAATATTGTCCC | This work |
| **Primers to create the Δ*rsh*::janus amplicon** | | | |
| RSG63 | ~1000bp upstream *rsh* | ACAGGATTCACGGTTTTATGG | This work |
| RSG64 | ~1000bp downstream *rsh* | CGTGCAGGATAGGATACCC | This work |
| RSG61 | End Janus cassette, **overlap just down *rsh*** | GTCCAAAAGCATAAGGAAAG**TTGTCCTAGCTCTTACTAGAAAG** | This work |
| RSG62 | Start Janus cassette, **overlap just up of *rsh*** | CACATTATCCATTAAAAATCAAAC**CTCTACTCTCCAATTCTTCCT** | This work |
| **Primers to create the *PcomX*::*alaRS*$_{editing}$ amplicon** | | | |
| KHB31 | ~800bp upstream *PcomX* | ATAACAAATCCAGTAGCTTTGG | (Berg et al., 2011) |
| KHB34 | ~800bp downstream *PcomX*::janus | CATCGGAACCTATACTCTTTTAG | (Berg et al., 2011) |
| RSG65 | end of PcomX, **overlap start *alaRS*$_{editing}$** | ATTTATATTTATTATTGGAGGTTCA**GCGTCAGCTGTCAAGGGTG** | This work |
| RSG66 | just downstream *PcomX*::janus, **overlapp end *alaRS*$_{editing}$** | ATTGGGAAGAGTTACATATTAGAAA**TTACAATTTACCTGCTACTGCATC** | This work |
| **Primers to introduce *murM*$^{K35A,R215A}$ and *murM*$^{K35A,Y219A}$** | | | |
| VE47 | ~1000bp upstream *murM*$^{R6}$ | ACCAGTAGTCATGGAAGCAAA | (Berg et al., 2013) |
| AW496 | Introduces the mutations K35A and W38A in MurM | **CGCATCAGAAGC**CACTTTTTCCCAAGCACTGC | This work |
| AW497 | Introduces the mutations K35A and W38A in MurM | **GAAAAAGTGGCTTCTGATGCG**AATCATGAGAGACTTGGTGTCTA | This work |
| AW498 | Introduces the mutations R215A and Y219A in MurM | **AGCAGCTTCGTTCGC**TAAATGAATCTCTTTTCGTTTCTCAG | This work |

| | | | |
|---|---|---|---|
| AW499 | Introduces the mutations R215A and Y219A in MurM | **CATTTAGCGAACGAAGCTGCT**TATAGAAAA TTATTAGATAACTTCAAAGAA | This work |
| MVH44 | ~100 bp downstream *murN*$^{R6}$ (also ~1300bp downstream *murM*$^{R6}$) | GCCAGCGCATGTCTCTCC | This work |
